# Redox regulation of neuroinflammatory pathways contributes to damage in Alzheimer’s disease brain

**DOI:** 10.1101/2025.09.08.674963

**Authors:** Lauren N. Carnevale, Piu Banerjee, Xu Zhang, Jazmin Navarro, Charlene K Raspur, Parth Patel, Tomohiro Nakamura, Emily Schahrer, Henry Scott, Nhi Lang, Jolene K. Diedrich, Amanda J. Roberts, John R. Yates, Stuart A. Lipton

## Abstract

The mechanism(s) whereby redox stress mediates aberrant immune signaling in age-related neurological disorders remains largely unknown. Normally, the innate immune system mounts a robust response to infectious stimuli. However, unintentional activation by host-derived factors, such as aggregated proteins associated with neurodegenerative disorders or by cytoplasmic genomic or mitochondrial DNA, can elicit aberrant immune responses. One such immune response is represented by the cytosolic GMP-AMP synthase–stimulator of interferon genes (cGAS-STING) pathway. Using redox chemical biology and mass spectrometry approaches, we identified *S*-nitrosylation of STING cysteine 148 as a novel posttranslational redox modification underlying aberrant type 1 interferon signaling in Alzheimer’s disease (AD). Critically, we observed *S*-nitrosylated STING (SNO-STING) in postmortem human AD brains, in hiPSC-derived microglia (hiMG) exposed to amyloid-β (Aβ)/α-synuclein (αSyn) aggregates, and in 5xFAD transgenic mice. Mechanistically, our findings reveal that STING *S*-nitrosylation is critical in initiating signaling cascades by promoting the formation of disulfide-bonded STING oligomers. This leads to neuroinflammation early in the course of disease *in vivo* in 5xFAD mice with consequent synaptic loss. Collectively, our research supports the role of SNO-STING in neuroinflammation associated with AD, and points to a novel druggable cysteine residue in STING to prevent this *S*-nitrosylation reaction with its inherent inflammatory response.

**One Sentence Summary:** *S*-Nitrosylation of STING triggers activation of cGAS-STING signaling in Alzheimer’s disease brain and subserves a novel link between excessive nitrosative stress and dysregulated innate immunity, thus contributing to disease progression.

## Introduction

The innate immune system protects the human body from disease by sensing infectious stimuli and coordinating a rapid yet controlled response but has also been implicated in the pathogenesis of neurodegenerative disorders such as Alzheimer’s disease^1,2^. Detection of foreign DNA is an essential component of innate immunity that is evolutionarily conserved across multiple species. The nucleic acid sensor cyclic GMP–AMP synthase (cGAS) detects cytoplasmic DNA (double-stranded DNA or DNA: RNA hybrids) and synthesizes 2,3-cyclic GMP-AMP (2,3-cGAMP). 2,3-cGAMP is a secondary messenger that activates the endoplasmic reticulum (ER) membrane protein stimulator of interferon genes (STING). Upon docking to STING, the cytosolic 2,3-cGAMP binding domain of STING undergoes a 180° rotation leading to the formation of adjacent STING dimers. After dimerizing, human STING is thought to polymerize by inter-dimer disulfide cross-linking mediated via cysteine 148, forming 2,3-cGAMP-bound STING oligomers that stabilize the active conformation. Moreover, some forms of oxidation of cysteine 148 are known to suppress STING signaling, most likely involving destabilization of the STING oligomers and dysregulation of oxidative stress pathways^3,4^ These oligomers are the active platform essential for inducing the expression of interferon-stimulated genes (ISGs). Therefore, hereafter we designate the active form as STING oligomers.

STING oligomers translocate to the Golgi apparatus and interact with TBK1. Critical interactions of the STING oligomers and TBK1 lead to a series of phosphorylation events. TBK1 undergoes autophosphorylation (p-TBK1) and mediates the phosphorylation of STING (p-STING) at serine 366, which leads to the TBK1-mediated phosphorylation of transcription factor IRF3 (p-IRF3). p-IRF3 then translocates to the nucleus and transcribes ISGs^5^ Although this immune surveillance can protect the body from infection, unintentional activation is detrimental^6^. For example, after mitochondrial damage, mislocalized mitochondrial DNA (mtDNA) accumulates in the cytosol in neurodegenerative diseases and drives neuroinflammation via ISGs as well as other inflammatory pathways such as the NLR family pyrin domain containing 3 (NLRP3) inflammasome^7–9^

Alzheimer’s disease (AD) is a neurodegenerative disease and is the most common form of dementia that occurs with aging. Neuropathological features of AD include the presence of extracellular amyloid-beta (Aβ) plaques and hyperphosphorylated tau in intraneuronal neurofibrillary tangles, often accompanied by accumulation of other aggregated proteins (e.g., α-synuclein (αSyn)), with resultant neuroinflammation, metabolic dysregulation, synapse loss, and consequent progressive cognitive decline^10–12^. A small set of familial AD cases have a clear genetic cause, but >95% cases are sporadic. Clinical and preclinical studies have identified nitrosative stress caused by reactive nitrogen species (RNS), such as nitric oxide (NO), as secondary signaling molecules driving age-related disease including AD^13,14^. Recent work also demonstrates a strong link between innate immune dysfunction and AD. For example, NLRP3 inflammasome activation and type 1 interferon signaling drive cellular damage, synaptic loss, and neuronal death^15–17^. Furthermore, Aβ pathology was observed to drive type 1 interferon activation in microglia and lead to memory and synaptic deficits, demonstrating that interferon signaling is a critical component of neuroinflammation in AD that is detrimental to memory and cognition^18^. Excessive NO, a known mediator of the immune response, induces nitrosative stress by modifying the dynamics of biological targets. Protein *S*-nitrosylation is a reversible redox-driven posttranslational modification (PTM) in which NO-related species covalently react with a cysteine thiol (or, more properly, thiolate anion) to form an S-nitrosothiol (SNO) adduct^19–22^. Protein *S*-nitrosylation diversifies proteome function without affecting protein expression, and regulates biological processes underlying health and disease. Here, we focus on identifying redox-regulated inflammatory pathways involved in the induction of type 1 interferons. Accordingly, we studied proteins involved in the initial “sensing” stages of innate immunity set the stage for downstream inflammation and cellular damage. Identifying such proteins and their redox-active cysteine(s) could open new opportunities for therapeutic intervention.

STING is involved in activating type 1 interferon and inflammasome signaling, both of which contribute to neuroinflammation^16,23,24^. STING is expressed in microglia, the innate immune cells of the central nervous system (CNS), and neurons, suggesting an interplay of type 1 interferon signaling in cell types relevant to AD. Furthermore, there are no STING-targeting therapeutics in clinical trials to date– an indication of the incomplete understanding of STING’s complex activation mechanisms. It is therefore essential to understand critical STING activation and regulatory events in order to successfully modulate neuroinflammation. In the present study, we identify protein *S*-nitrosylation (SNO) as a novel and critical STING PTM in human induced pluripotent stem cell (hiPSC)-derived microglia (hiMG) and human monocytes (hM), and also in human brain tissue from patients with AD. Our findings indicate that SNO is necessary for effective cGAS-STING signaling, whereas inhibition of this reaction suppresses STING oligomerization and the production of interferon-β (IFN-β). We demonstrate altered levels of cGAS-STING signaling proteins, including an increased p-TBK1/TBK1 ratio in male and female sporadic AD brains compared to healthy brains. Our data illuminate *S*-nitrosylation of human STING as a key redox-mediated activation step underlying neuroinflammatory damage in AD.

## Results

### Dysregulated cGAS-STING signaling and presence of SNO-STING in human postmortem Alzheimer’s disease brain tissue

We characterized the expression of a panel of cGAS-STING pathway proteins in human postmortem brains from healthy controls and sporadic AD patients of both sexes, and found that STING was *S*-nitrosylated (**Fig. 1a**). SNO-STING was significantly increased in human AD brains relative to controls succumbing to non-CNS diseases. We next investigated cGAS-STING-dependent downstream signaling to better understand the implications of STING *S*-nitrosylation. In this regard, we observed a significant increase in the ratio of p-TBK1/TBK1 in both female and male samples (**Fig. 1b-e**). Interestingly, only select female AD brain samples had statistically significant levels of elevated p-IRF3. Furthermore, by examining the subset of patient samples for which the ApoE status was known, we observed that female AD brains containing the AD-risk allele, ApoE4, had the greatest number of cGAS-STING markers altered compared to control female brains (**Table S1**). These findings may be of particular importance given the increased vulnerability of women to AD^25^

**Fig. 1.**
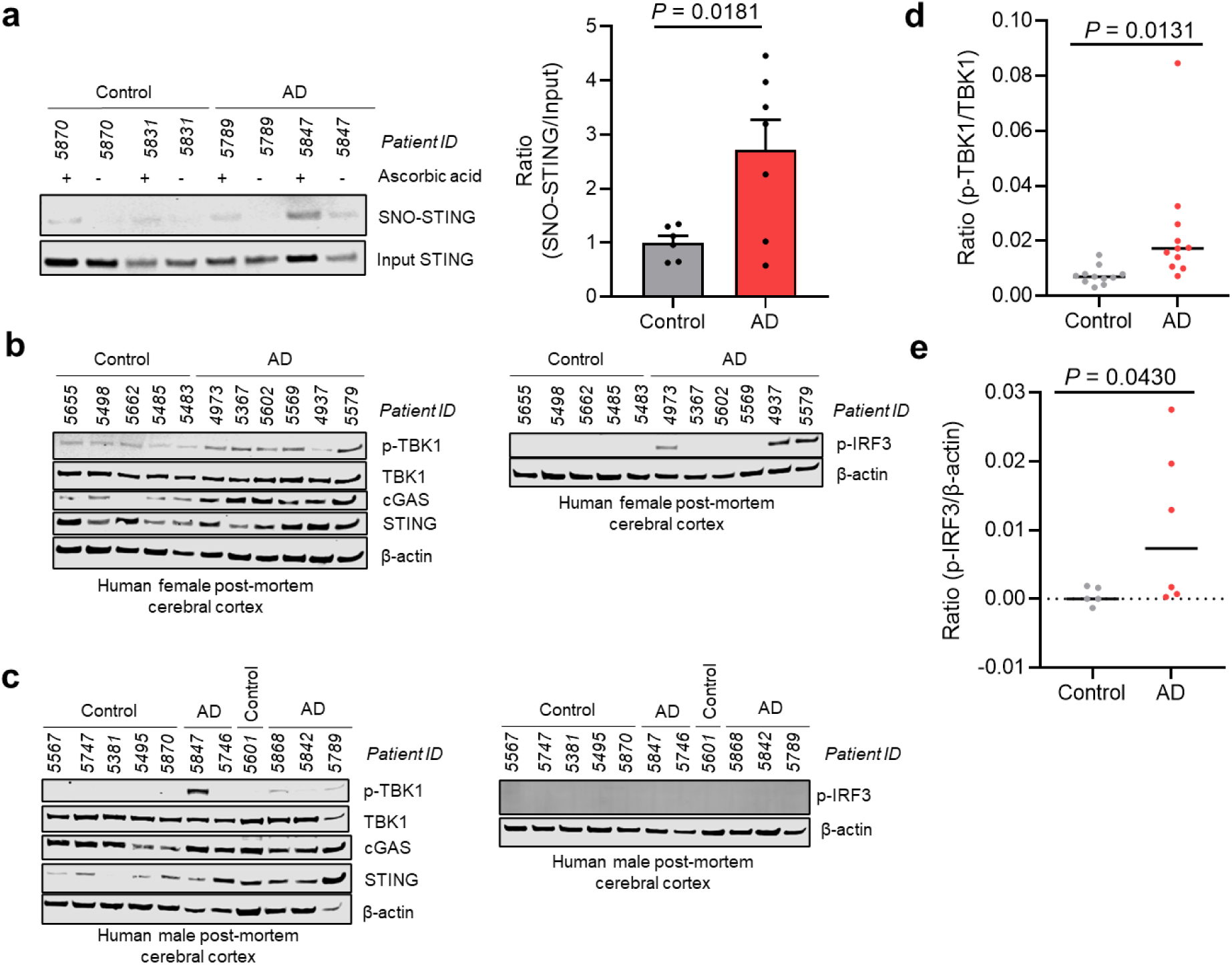
Human postmortem AD brain contains *S*-nitrosylated STING and active cGAS-STING signaling. **a,** Detection of SNO-STING in control (*n* = 6) and AD (*n* = 7) postmortem human brain lysates. Bar graph shows mean + s.e.m. of relative ratio of SNO-STING to total STING. **b,c,** Detection of p-TBK1, cGAS, STING, and p-IRF3 in male (**b**) and female (**c**) control and AD postmortem brain lysates. Proteins were detected using SDS-PAGE and immunoblotting. **d,e,** Quantification of protein levels AD brain compared to control for p-TBK1/TBK1 and cGAS/ β-actin. Data are individual values and mean (horizontal line). *P* values determined by Student’s *t* test.

### *S*-Nitrosylation of STING in human immune cells

Microglia play an essential role in the pathogenesis of AD by contributing to neuroinflammation and are known to accumulate around Aβ plaques^26–28^. We confirmed the presence of cGAS-STING protein machinery in our hiMG culture system and their functional response to 2,3-cGAMP treatment. We observed the formation of STING oligomers in response to 2,3-cGAMP treatment (**Extended Data Fig. 1).** We then investigated whether *S*-nitrosylation of STING occurs in hiMG and hM (**Fig. 2**). Cells were exposed to the physiological NO donor/transnitrosylating agent S-nitrosocysteine (SNOC) at room temperature, and after 20 min, a biotin-switch assay was performed. SNOC was observed to induce SNO-STING formation in hiMG and hM (**Fig. 2b,c**). Notably, SNOC has a very short half-life, on the order of 30 seconds to a minute at physiological pH and room temperature, so this reaction must have occurred relatively rapidly.

**Fig. 2.**
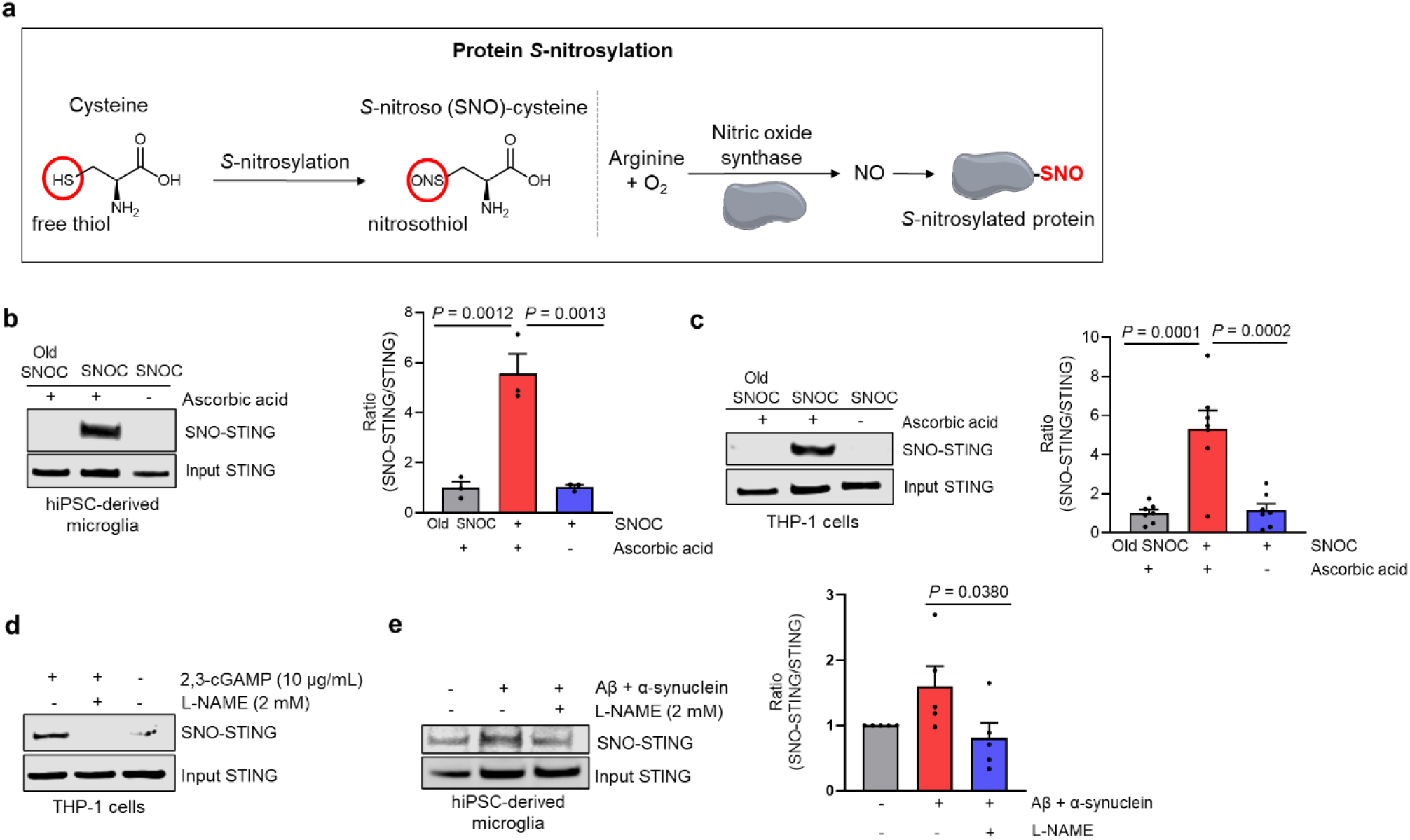
STING is *S*-nitrosylated in immune cells. **a,** Simplified schema of protein *S*-nitrosylation. **b,** *S*-Nitrosylation of human STING in hiMG. Cells were exposed to the physiological NO donor/transnitrosylating agent S-nitrosocysteine (SNOC, 200 μM) at room temperature. Biotin switch was performed 20 minutes later to assess S-nitrosylated STING (SNO-STING). This method detects protein *S*-nitrosylation through sequential blocking of free thiols, switching the *S*-nitrosothiol to a biotin label following ascorbate reduction, and, finally, detection of the biotinylated protein. Samples without ascorbate addition (ascorbic acid -) or old SNOC (from which NO had been dissipated) served as negative controls. Data represent mean s.e.m. (*n* = 3 experiments). Statistics by ANOVA and Tukey’s multiple comparison’s test. **c,** SNO-STING formation in hM (THP-1 cells). Data represent mean + s.e.m. (*n* = 7 experiments). Statistics by ANOVA and Tukey’s multiple comparison’s test. **d,** *S*-Nitrosylation of STING triggered by its endogenous ligand 2,3-cGAMP. SNO-STING formation was abrogated in cells pre-incubated in NOS inhibitor L-NAME for 3 h prior to addition of 2,3-cGAMP for 24 h. **e,** Aβ and αSyn oligomers induce SNO-STING formation in hiMG. Cells were exposed to protein oligomers (Aβ, 750 nM; αSyn, 150 nM) for 24 h prior to SNO-STING detection by biotin-switch assay. Data represent mean + s.e.m. (*n* = 5 experiments). Statistics by a Student’s unpaired *t* test.

Activation of STING by the cGAS-derived secondary messenger, 2,3-cGAMP, leads to downstream type 1 interferon signaling. Therefore, we evaluated whether 2,3-cGAMP contributes to *S*-nitrosylation of STING. We observed that 2,3-cGAMP led to SNO-STING formation and that inhibiting NO production with the NO synthase (NOS) inhibitor, L-N^G^-nitroarginine methyl ester (L-NAME), abrogated this reaction (**Fig. 2d**).

Soluble extracellular aggregates of Aβ_1-42_ and αSyn are thought to play a role in the progression and pathogenesis of AD^29^. Moreover, exposure of hiMG to Aβ_1-42_ and αSyn oligomers causes activation of the NLRP3 inflammasome^29^, and leads to neuronal synapse loss and eventual cell death^30,31,40^. Interestingly, we found that hiMG exposed to Aβ_1-42_ and αSyn oligomers demonstrated increased SNO-STING, an effect also prevented by L-NAME (**Fig. 2e**). These results suggest that Aβ_1-42_ and αSyn oligomers may contribute to aberrant innate immune signaling via *S*-nitrosylation of STING. Moreover, after exposure of hiMG or hM to Aβ_1-42_ and αSyn oligomers, we observed an increase in the ratio of p-TBK1/total TBK1 as well as increased interferon signaling (**Extended Data Fig. 2**). These findings are consistent with the notion that AD-relevant protein aggregates trigger cGAS-STING inflammatory signaling that contributes to the neurodegenerative process.

Mechanistically, endogenous STING agonist 2,3-cGAMP induces a closed conformation of activated STING through the formation of a lid region and a 180° rotation of the ligand-binding domain. In this state, 2,3-cGAMP is buried within the dimer interface^32^. We found that co-exposure of either human or mouse monocytoid cells to 2,3-cGAMP plus the NO donor SNOC elevated the ratio of p-TBK1/TBK1 and relative IFN-β mRNA levels above that of 2,3-cGAMP alone, supporting the notion that *S*-nitrosylation is a critical STING activation step triggering downstream signaling (**Extended Data Fig. 3**).

### *S*-Nitrosylation of STING occurs predominantly at cysteine 148

To identify the specific STING cysteine residue involved in *S*-nitrosylation, we performed mass spectrometry on human recombinant STING after exposure to SNOC. The results showed that cysteine 148, a critical residue known to mediate structural changes in STING oligomerization and activation, is *S*-nitrosylated (**Fig. 3a**). We next overexpressed wild-type (WT) FLAG-tagged STING or cysteine-to-alanine (C148A) mutant and performed biotin-switch assays. After exposure to SNOC, only STING^WT^ SNOC exhibited a significant increase in *S*-nitrosylation and oligomerization, confirming that cysteine 148 is responsible for STING protein *S*-nitrosylation (**Fig. 3b,c**).

**Fig. 3.**
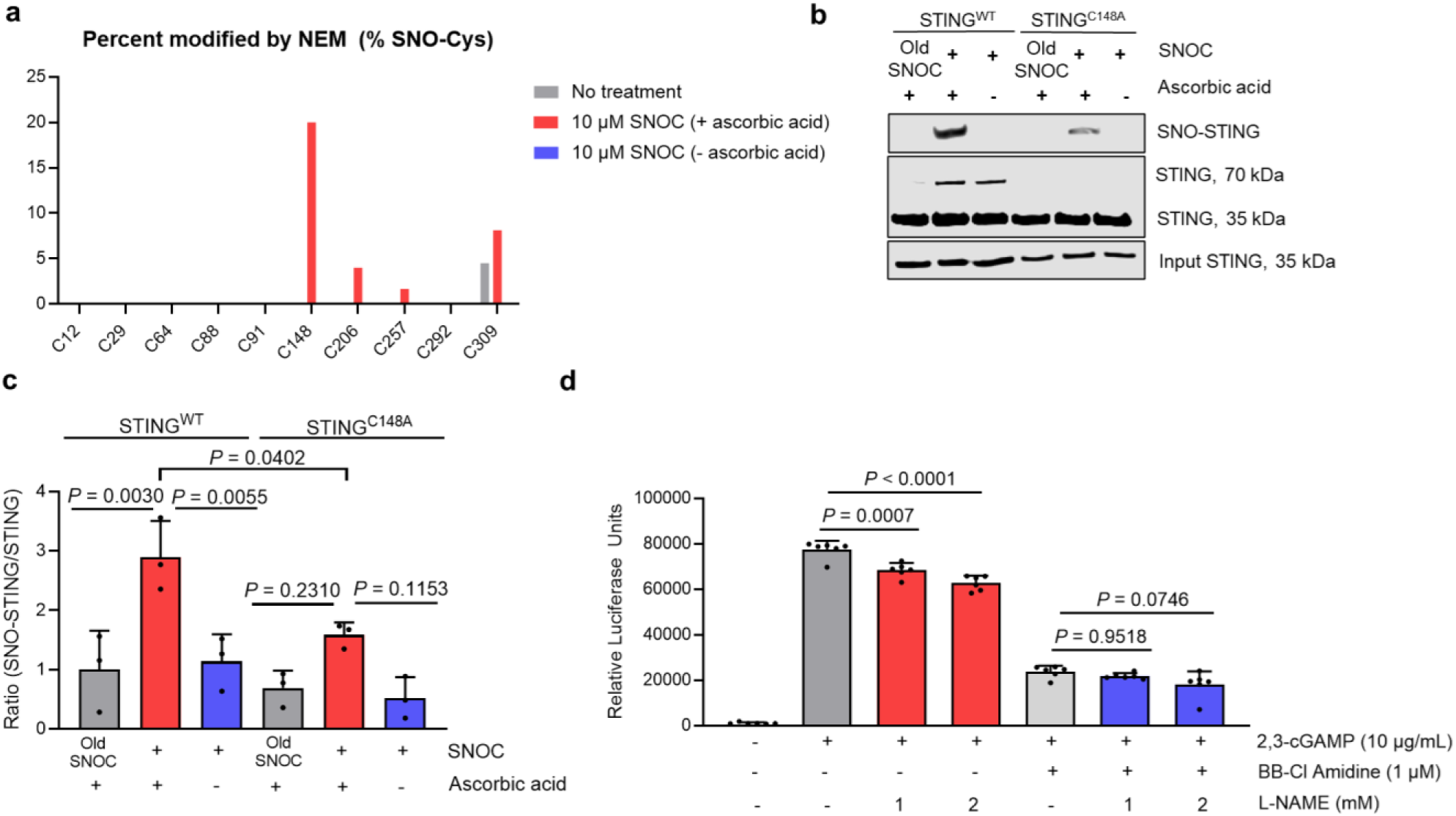
Cysteine 148 undergoes *S-*nitrosylation and leads to NO-dependent disulfide-linked STING species. **a,** Detection of *S-*nitrosylated cysteine residues using mass spectrometry. **b,** Detection of SNO-STING by the biotin-switch assay (*n* = 3). HEK 293T cells were transfected with WT or C148A human STING-EGFP. **c,** Quantification of biotin-switch assays. Values are mean + s.e.m., ANOVA with Tukey’s multiple comparison’s test (*n* = 3). **d,**THP-1 Dual hM were pre-treated with NOS inhibitor L-NAME for 3 h. During the final hour of pre-treatment, the STING inhibitor BB-Cl Amidine was added to the cells. Cells were then stimulated with 2,3-cGAMP, and 24 h later interferon signaling was assessed by measuring luciferase activity. Values are mean + s.e.m., ANOVA with Tukey’s multiple comparison’s test (*n* = 6).

To further investigate the role of cysteine 148 on the NO-dependence of cGAS-STING signaling, we utilized the covalent STING inhibitor BB-Cl Amidine, which specifically modifies STING at cysteine 148. THP-1 Dual hM were pre-treated with L-NAME for 3 hours, with BB-Cl Amidine added during the final hour. Following stimulation with 2,3-cGAMP for 24 h, we found that covalent modification of cysteine 148 effectively inhibited NO-dependence on STING signaling (**Fig. 3d**). Prior to this work, BB-Cl Amidine was not known to modify a cysteine residue involved in protein *S*-nitrosylation. This finding highlights the therapeutic potential of targeting a cysteine residue involved in *S*-nitrosothiol formation to fine tune immune signaling. The efficacy in preventing STING signaling through covalent modification of cysteine 148 specifically demonstrates one such candidate cysteine.

### *S*-Nitrosylation mediates STING oligomerization

Docking of 2,3-cGAMP at STING’s binding pocket induces a conformational change, leading to transient STING oligomerization. Current evidence suggests that STING oligomerization triggers downstream type 1 interferon signaling^3^. When we treated hM with 2,3-cGAMP (10 μg/mL), we observed the disappearance of STING monomers and the appearance of oligomers within 1 h, after which the ratio of oligomers to monomers decreased (**Fig. 4a-b**). Next, we tested whether STING oligomers induced by 2,3-cGAMP is dependent on endogenous NO. Under conditions that generated SNO-STING, we incubated THP-1 Dual hM (containing a luciferase reporter for ISGs) with 2,3-cGAMP (10 μg/mL) in the presence or absence of _L_-NAME for 24 h. We observed a decrease in ISG signaling in cells treated with L-NAME consistent with the notion that endogenous NO regulates the activity of this pathway (**Fig. 4c, Extended Data Fig. 3**). We also observed SNO-STING and STING oligomer formation in the presence of 2,3-cGAMP or lipopolysaccharide (LPS) in hM, which was dependent on endogenous NO (**Fig. 4d,e**). In hM exposed to increasing concentrations of SNOC, we observed the parallel formation of SNO-STING and STING oligomers (**Fig. 4f**). Furthermore, when we incubated full-length recombinant human STING with increasing concentrations of the SNOC, we observed formation of STING oligomers in a concentration-dependent manner (**Fig. 4g**). These results suggest that NO facilitates STING oligomerization. As additional support for this premise, the synthetic STING agonist SR-717, which adopts a closed-state STING conformation like 2,3-cGAMP,^33^ also led to the formation of NO-dependent STING oligomers (**Fig. 4h**). Taken together, these findings indicate that both endogenous and synthetic agonists rely on cellular NO levels for STING oligomerization/activation. Accordingly, we propose a model in which *S*-nitrosylation facilitates STING oligomerization and thus contributes to downstream type 1 interferon signaling (**Fig. 4i**).

**Fig. 4.**
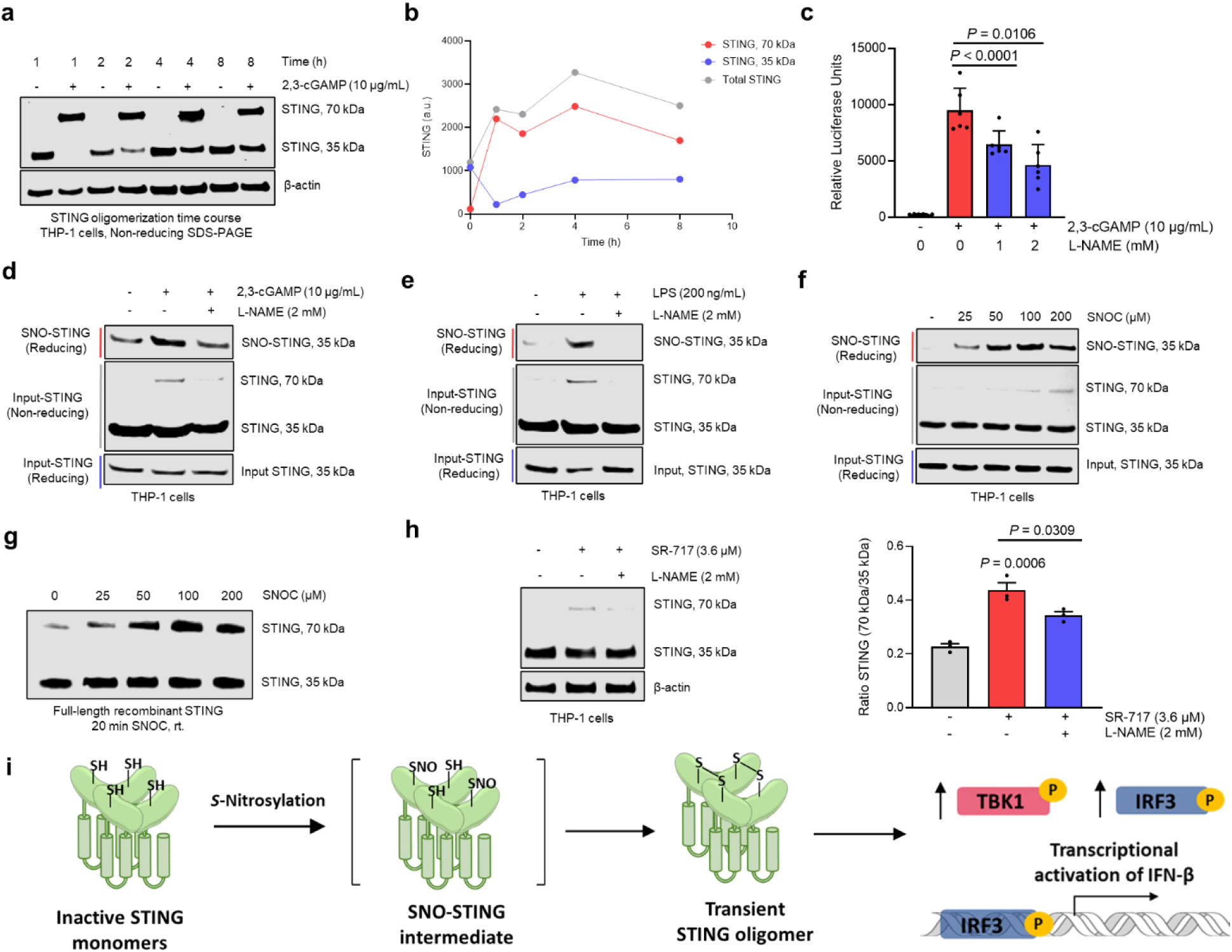
*S*-Nitrosylation mediates STING oligomerization in hM. **a,b,** 2,3-cGAMP induced STING oligomerization, usually observed as dimers after gel electrophoresis.^51^ hM were treated with 2,3-cGAMP (Invivogen) for 1-8 h. Non-reducing SDS-PAGE was used to observe STING monomers and oligomers. a.u., arbitrary units. **c,** THP-1 dual hM were pre-treated with L-NAME for 3 hours before treatment with 2,3-cGAMP. Ordinate axis shows interferon signaling monitored by luciferase activity after 24 h. Values are mean + s.e.m., ANOVA with Tukey’s multiple comparison’s test (*n* = 6). **d,e,** hM cells were pre-incubated without or with L-NAME for 3 h before treatment with 2,3-cGAMP (**d**) or LPS (**e**) for 24 h. SNO-STING and STING oligomers were observed using reducing and non-reducing SDS-PAGE, respectively. **f,** hM cells were exposed to SNOC (25-200 μM) at room temperature, followed 20 min later by analysis of SNO-STING and STING oligomerization. **g,** Full-length recombinant human STING was exposed to SNOC at room temperature. Non-reducing SDS-PAGE was performed 20 minutes later to observe STING oligomerization. **h,** Synthetic closed-state STING agonist SR-717 induced STING oligomers that were abrogated by L-NAME (*left*). Values are mean + s.e.m. (*n* = 3), ANOVA with Tukey’s multiple comparison’s test (*right*). **i,** Schematic of proposed *S*-nitrosylation mechanism underlying STING oligomerization. STING dimerizes and then oligomerizes, facilitated by S-nitrosylation-induced disulfide bond formation, leading to phosphorylation of TBK1 and IRF3 with downstream type 1 interferon activation.

We then utilized a second NO donor, spermine NONOate, to further assess STING oligomerization. Spermine NONOate is a chemical tool developed for the slow release of NO. It follows first-order kinetics with a half-life of 39 minutes at 37 °C. hM cells exposed to spermine NONOate (4 – 400 μM) for 3 h led to STING oligomerization (**Extended Data Fig. 4**). As spermine can condense viral DNA, but not host nucleosome DNA, leading to cGAS activation, we included a spermine control to investigate whether spermine alone (without NO) could initiate STING oligomerization^34^, but as a negative control, we did not observe STING oligomers in cells exposed to equimolar spermine alone. This indicates that NO-related species, but not spermine, led to the formation of STING oligomers (**Extended Data Fig. 4**). Given that *S*-nitrosylation of a member of a vicinal pair of cysteines is known to facilitate disulfide bond formation^35^ and that human STING is thought to polymerize via inter-dimer disulfide formation at cysteine 148, the very site we found to be SNOed on STING, it likely that formation of SNO-STING facilitates its oligomerization via disulfide formation.

### *S*-Nitrosylation stimulates interferon signaling by boosting STING activation

Next, we investigated whether NOS inhibitors could decrease relative mRNA expression of IFN-β. We found that L-NAME decreases IFN-β mRNA levels induced by 2,3-cGAMP in hM (**Extended Data Fig. 3)**. We then investigated whether brief exposure to SNOC affects 2,3-cGAMP-induced type 1 interferon signaling. We reasoned that if *S*-nitrosylation of STING facilitates it oligomerization in conjunction with 2,3-cGAMP binding, we would expect to see an increase in cGAS-STING signaling with the addition of NO, for example, via SNOC exposure. In fact, even if SNOC exposure preceded 2,3-cGAMP treatment, this enhancement in STING signaling should be observed since S-nitrosylation of cysteine 148 would occur prior to binding of 2,3-cGAMP but would subsequently facilitate disulfide formation when vicinal cysteine residues approached each other, as known to occur in the presence of 2,3-cGAMP. To explore this possibility, we pre-incubated BV2 mouse microglia with SNOC, and 10 min later replaced the media with fresh solution containing 2,3-cGAMP for 60 min. We found that under these conditions SNOC boosted 2,3-cGAMP-induced STING oligomerization, p-TBK1 levels, and relative IFN-β mRNA levels (**Extended Data Fig. 3**). SNOC alone did not lead to elevated IFN-β mRNA expression, suggesting that STING oligomerization alone is insufficient to elicit the transcriptional activation of IFN-β.

Based on these observations, STING may require two ‘hits’ for activation: (i) an *S*-nitrosylation event to facilitate subsequent oligomerization, and (ii) 2,3-cGAMP to induce a structural change in STING, e.g., dimerization, so that subsequent disulfide bond formation can occur, producing an increasing number of dimers or dimers, leading to polymerization. Accordingly, we found that that co-incubation with 2,3-cGAMP and NO donor boosted cGAS-STING signaling beyond that of 2,3-cGAMP alone (**Extended Data Fig. 3**). In fact, the effect of NO in stimulating STING activity was dependent on the presence of 2,3-cGAMP, suggesting a permissive action of 2,3-cGAMP, consistent with the mechanistic explanation offered above (**Fig. 4i**).

### *S*-Nitrosylation regulates STING condensates

Protein biocondensates are non-membranous cellular compartments that arise from liquid-liquid phase separation (LLPS). LLPS is a biophysical process driven by soluble proteins containing weak multivalent interactions, modular domains, or intrinsically disordered regions. The LLPS of several proteins involved in neurodegeneration and antiviral signaling have been characterized, but their regulatory mechanisms are unclear^36,37^. Biocondensates create a boundary from the environment that limit molecular exchange, and alter biological kinetics and local protein concentrations, impacting biological processes in diverse ways. STING biocondensates, which can be observed as puncta, form in the presence of 2,3-cGAMP and function as a regulatory mechanism to prevent hyperactivation^37^. As PTMs modulate protein function without changing protein abundance, we sought to investigate the effect of protein *S*-nitrosylation on STING biocondensates. Based on our data indicating that NO promotes SNO-STING formation and type 1 interferon signaling, we hypothesized that NO might also limit STING biocondensates, thus disrupting their regulatory role, and thereby contributing to STING hyperactivation. Consistent with prior studies, we observed STING biocondensates in cells overexpressing STING and an increased number of STING biocondensates in the presence of 2,3-cGAMP. Moreover, both of the NO donors used here, SNOC and spermine NONOate, decreased formation of STING biocondensates in the presence or absence of 2,3-cGAMP (**Extended Data Fig. 5**). Intriguingly, this finding is consistent with prior studies that utilized cells containing STING mutations underlying STING-associated vasculopathy with onset in infancy (SAVI), an autoimmune disease characterized by chronic hyperactivation of type 1 interferon signaling. SAVI mutations, including V147L, N154S, and V155M, occur at the STING polymer interface near cysteine 148 (C148), the site we found to be S-nitrosylated^38,39^. Similar to our findings in cells treated with NO donors, the previous studies found that cells with SAVI mutations manifest fewer STING biocondensates, thereby disrupting a key regulatory event and contributing to hyperactivation and prolonged inflammation. We speculate that the structure of SNO-STING may mimic the known genetic STING mutations associated with SAVI, leading to this hyperactivated STING state^37^. We also demonstrated that 2,3-cGAMP does not increase STING levels (**Fig. 2d and Fig. 4d**); and neither does SNOC nor spermine NONOate (**Fig. 2b,c and Extended Data Fig. 4**). Therefore, it is unlikely that NO inhibited STING biocondensates due to a decrease in STING protein levels. Rather, it would appear that the ordered array imposed by protein *S*-nitrosylation-induced oligomerization of STING via polymer formation favors the active conformation and avoids LLPS.

### Presence of SNO-STING in 5xFAD transgenic mice

Similar to human AD brain, recent work has demonstrated that the 5xFAD mouse brain exhibits elevated cGAS-STING signaling.^40^ Therefore, we tested whether SNO-STING is present in 5xFAD mouse brains. Using age- and sex-matched, littermate WT vs. 5xFAD mice, we determined the levels of SNO-STING relative to input STING in each sample with and without ascorbate as a negative control. By 3 months of age, we observed an increased ratio of SNO-STING to total STING in 5xFAD mice (**Fig. 5**) compared to WT littermate mice. This finding is consistent with the notion that dysregulated cGAS-STING machinery occurs at an early stage of the disease since the 5xFAD mouse at this time point is just beginning to manifest behavioral, electrophysiological, and histological evidence of disease^41^

**Fig. 5.**
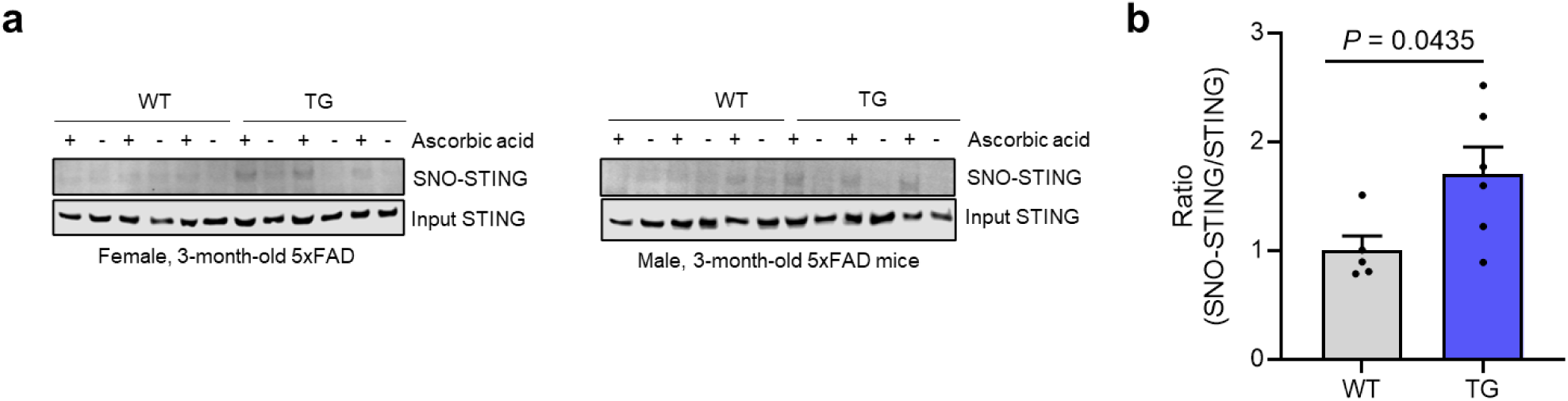
*S*-Nitrosylated STING in 3-month-old 5xFAD mice. **a,** Detection of SNO-STING in 3-month-old 5xFAD transgenic (TG) and WT (*n* = 6) mice. **b,** Quantification of SNO-STING in TG (*n* = 5) and WT (*n* = 6), Student’s *t* test with *P* value indicated.

To determine whether accumulation of SNO-STING in our AD model systems was of pathophysiological significance compared to postmortem human AD brain, we calculated the relative ratio of SNO-STING (by biotin-switch assay) to total STING (as quantified from immunoblots), using methods previously described^31,42,43^. In fact, in human AD brain this ratio was generally comparable to or greater than that encountered in both our hiMG cell-based models and in the brains of the 5xFAD mice (cf. **Fig. 1a**, **Fig. 2e, and Fig. 5b**), showing that our model systems manifest pathophysiologically relevant levels of SNO-STING. Furthermore, as the mice reached 6 months of age, increased levels of the inflammatory markers GFAP (for astrocytes) and Iba1 (for microglia) were observed in the 5xFAD mice, indicating that the inflammation persists (**Supplemental Data Fig. 1**).

### Reversal of neuroinflammatory signaling by non-nitrosylatable cysteine mutant STING in AD transgenic mice *in vivo*

To evaluate the role of *S*-nitrosylation of STING(C148) in regulating neuroinflammation in AD, we injected lentiviral constructs encoding either non-nitrosylatable mutant STING^C148A^ or WT (STING^WT^) into the hippocampus of 5-month-old 5xFAD mice. One month later, we assessed microglial activation by employing immunofluorescence staining for Iba1 using quantitative confocal microscopy (**Fig. 6a,b**). We observed significant differences in Iba1 expression between the STING^WT^ and STING^C148A^ groups, with 5xFAD mice injected with the STING^C148A^ lentiviral construct exhibiting a marked reduction in Iba1 staining compared to the STING^WT^- injected group. Quantification of Iba1-positive microglia in regions containing tdTomato (indicating successful lentiviral expression) displayed a significant decrease in Iba1 integrated density in the hippocampus of the STING^C148A^-injected 5xFAD mice (**Fig. 6c**). This decrease in Iba1 expression is consistent with the notion that the C148A mutation, which prevents *S*- nitrosylation, attenuates microglial activation via attenuation of STING activation. Note that cGAS-STING signaling can be non-cell autonomous^44^. Hence, it is likely that lentiviral vectors expressed in both glia and neurons affected cGAS-STING signaling in this paradigm.

**Fig. 6.**
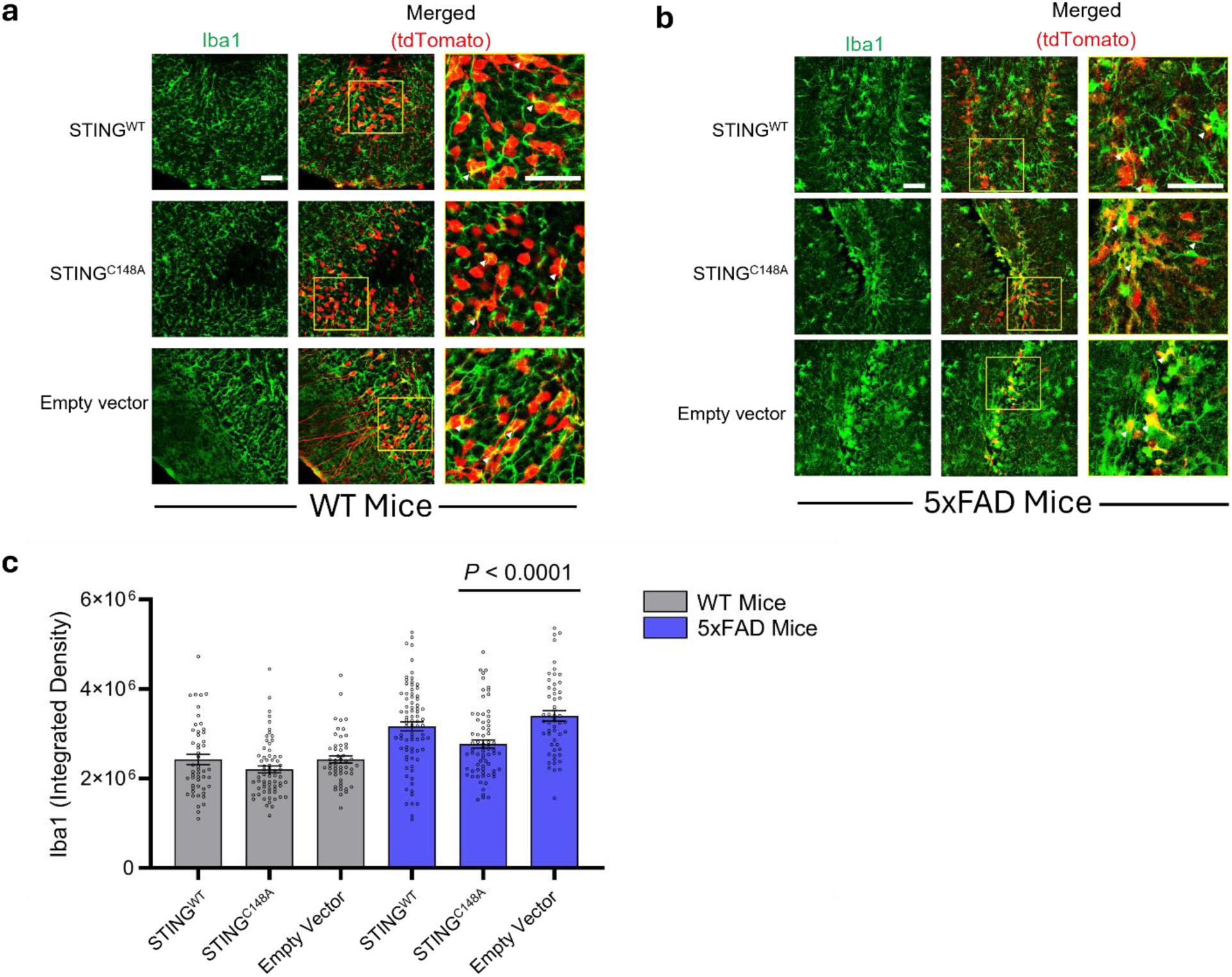
Lentiviral expression of non-nitrosylatable mutant STING^C148A^ *in vivo* decreases inflammation in 5xFAD mouse hippocampus. **a,b,** Representative images of hippocampal sections from WT mice **(a)** and 5xFAD mice **(b)** injected with either the STING^WT^, non-nitrosylatable mutant STING^C148A^, or empty vector, each labeled with tdTomato. Arrowheads indicate exemplary regions used for quantification expressing Iba1 (green) in the vicinity of tdTomato (red). Scale bars, 50 μm. **c,** Quantification of Iba1 integrated density in WT and 5xFAD mouse brains injected with each of the three vectors (*n* ≥ 50 for each group). Each datapoint represents an imaged region of interest (ROI). Bars show mean value ± s.e.m. Microglia in the region of expression of STING^C148A^ vector exhibited a significant decrease in integrated density compared to those expressing STING^WT^ or empty vector. Statistical significance by Student’s *t* test.

**Fig. 7.**
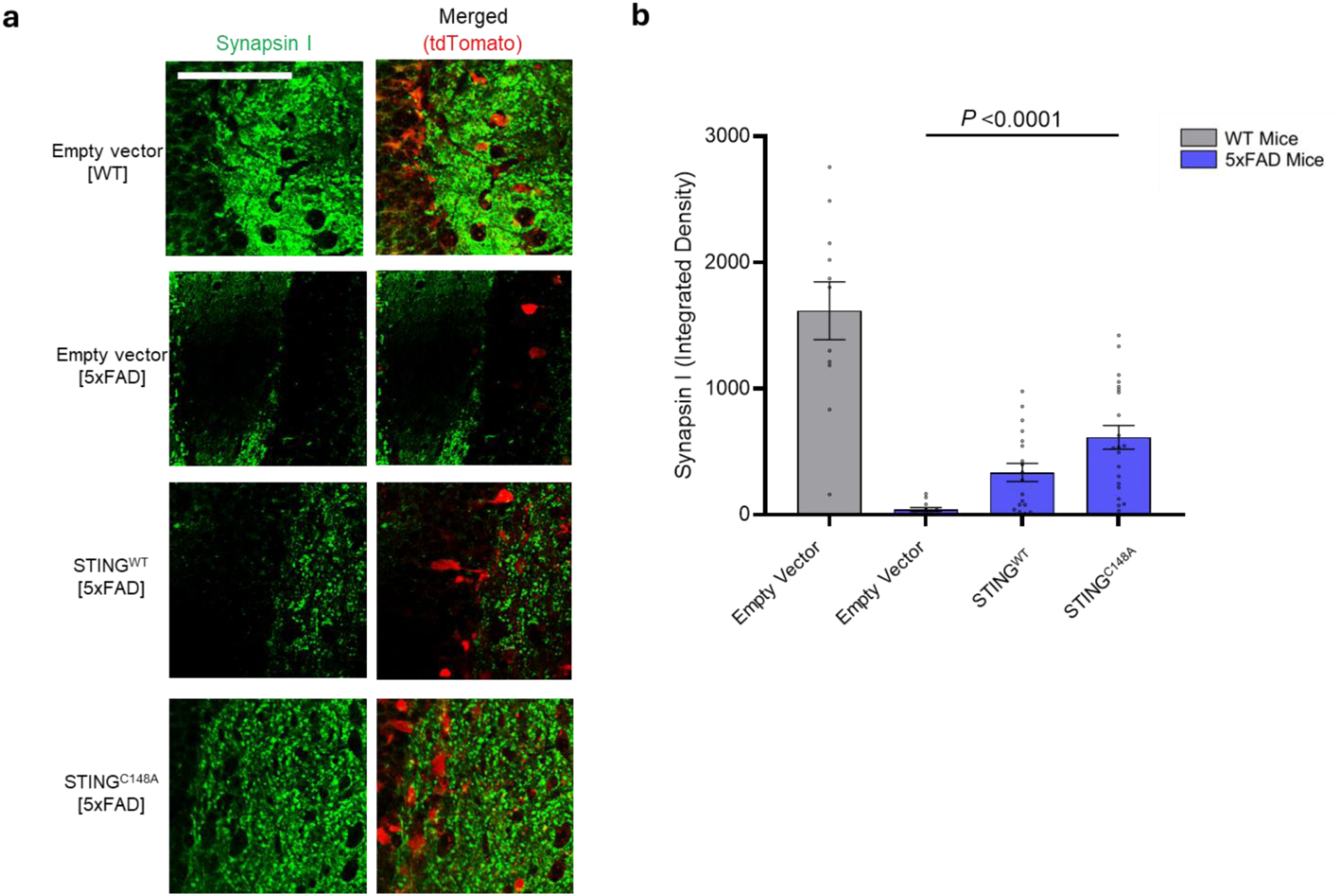
Lentiviral expression of non-nitrosylatable mutant STING STING^C148A^ *in vivo* partially protects synapses in 5xFAD mouse brain. **a,** Representative images of hippocampal sections from WT and 5xFAD mice injected with either STING^WT^, non-nitrosylatable mutant STING^C148A^, or empty vector, each labeled with tdTomato. Immunohistochemistry was performed with anti-Synapsin I antibody (green) to label synaptic boutons and anti-tdTomato antibody (red) to identify cells expressing the injected vector. Scale bar, 100 μm. **b,** Quantification of Synapsin I integrated density per unit area of ROI in WT mice (*n* = 11) and in 5xFAD mice injected with each of the three vectors (*n* ≥ 13 for each group). Each datapoint represents an individual ROI selected by the presence of tdTomato staining. Bars show mean value ± s.e.m. Statistical significance by Mann-Whitney U-test.

Additionally, we monitored a biomarker for synapse loss in the 5xFAD mouse brain as this represents the best clinical correlate to cognitive decline in human AD and has been linked to aberrant S-nitrosylation reactions in both microglia and neurons^25,30,31,45,46^. To determine specifically if *S-*nitrosylation of STING might contribute to synapse loss, we performed immunohistochemistry for Synapsin I, a marker of synaptic boutons, in areas of lentiviral expression of our STING constructs, as evidenced by tdTomato staining. Compared to lentiviral injection with empty vector, injection with non-nitrosylatable mutant STING^C148^, but not with STING^WT^, increased Synapsin I staining in a statistically significant manner, consistent with rescue from synaptic loss (**Fig. 8a,b**). Although there was also a trend toward improvement with STING^WT^, this may reflect overexpression of WT protein, which can act as a sink for NO, allowing substantial WT protein to remain un-nitrosylated, and therefore potentially offer some benefit, an effect we have reported previously in similar *in vivo* systems^31^. However, non-nitrosylatable mutant protein was more effective and showed statistically significant synaptic protection. Collectively, our findings indicate that non-nitrosylatable mutant STING^C148A^ can decrease neuroinflammation and synaptic loss in this *in vivo* transgenic mouse model of AD.

## Discussion

Current therapies for AD do not address the prominent neuroinflammatory component of the disease. In this regard, immune system dysregulation has been tied to the etiology of AD^23,40^, and the interplay of microglia and neurons is critical. The role of cGAS-STING signaling and, more specifically, STING activation has heretofore remained unclear in AD pathogenesis. In the current study, we provide evidence that postmortem human AD brain contains higher levels of *S-* nitrosylated STING, which contribute to activation of the cGAS-STING pathway. Our *in vitro* results demonstrate that NO-related species increase formation of STING oligomers and downstream interferon-stimulated genes, such as IFN-β. Importantly, we observed a similar pattern in human AD brains compared to controls.

Understanding the mechanism(s) underlying STING activation is essential because STING is involved in autophagy-mediated pathways, inflammasome activation, and type 1 interferon signaling, all thought to contribute to AD pathology. Accordingly, knowing the mechanism of STING activation should contribute to the development novel therapeutics^24,47^ Selectively modulating innate immunity in the CNS without impairing immune surveillance is key to preserving innate immunity in the CNS for protection against infectious processes^48–50^. Specifically, abrogating S-nitrosylation of STING in response to AD-related stimuli such as Aβ oligomers may afford the opportunity for such targeted intervention without altering the homeostatic functions of STING, such as autophagy.

Cysteine 148 on STING, the site we found to be S-nitrosylated, participates in disulfide-bond formation. Our findings suggest that SNO-STING facilitates the formation of this disulfide, which then contribute to formation of STING oligomers, essential for active cGAS-STING signaling. This finding suggests that the future therapy with STING inhibitors targeting cysteine 148 might be highly effective in abrogating excessive inflammatory signaling, as we show that protein *S*-nitrosylation of this critical residue, in conjunction with 2,3-cGAMP, acts as a key switch for oligomerization and pro-inflammatory interferon signaling. Moreover, cGAS-STING signaling itself may increase in NO levels, as NF-κB activation by cGAS-STING is upstream of NO production, thus creating a feed-forward effect on type 1 interferon signaling that would be modulated by the formation of SNO-STING. Interestingly, we also demonstrate that NO donors prevent formation of STING biocondensates. Previous work had suggested that STING biocondensates are a mechanism to prevent hyperactivation of cGAS-STING signaling^37^ Taken together with additional prior data^38,39^ showing that mutations at the polymer/oligomer interface, similar to the effect we see with S-nitrosylation, disrupt condensation of STING, our results are consistent with the notion that oligomerization of STING into active complexes also prevents its inactivation by LLPS into biocondensates.

In conclusion, our findings reveal STING as a redox-regulated protein that links nitrosative stress to neuroinflammation in AD brain and introduces *S-*nitrosylation as an important PTM that functionally activates STING without altering protein abundance. This discovery also affords the opportunity to target the site of S-nitrosylation, cysteine 148, therapeutically as an anti-inflammatory intervention for AD.

## Extended Figures and Extended Figure Legends

**Extended Data Fig. 1.**
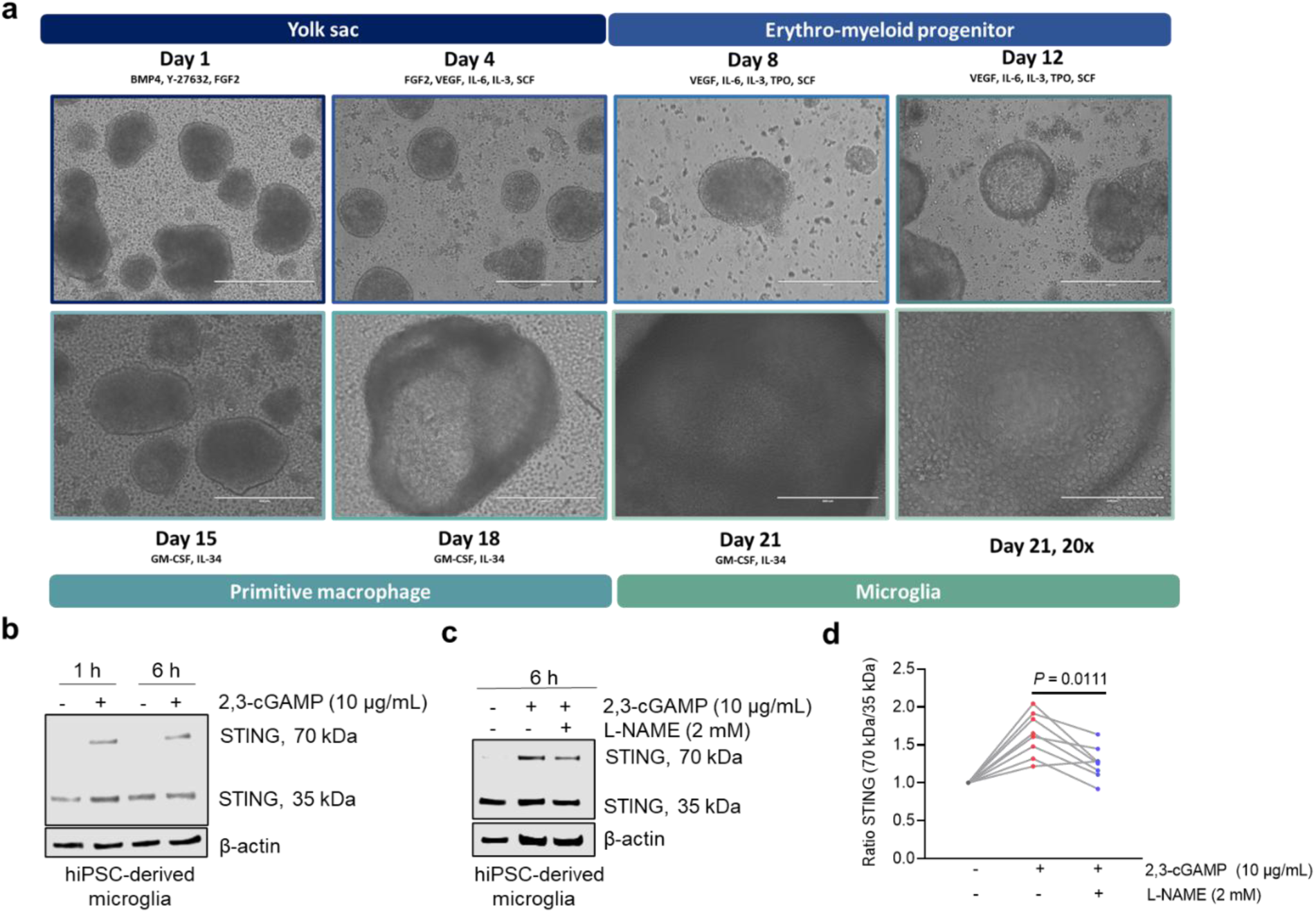
hiMG respond to 2,3-cGAMP. **a,** hiMG differentiation protocol. This timeline includes media supplements that drive differentiation of human iPSCs into microglia-like cells.^29^ Colored labels indicate each stage of the differentiation procedure along with day of differentiation. Scale bar, 400 μm; except day 21, 200 μm). **b,** Formation of STING oligomers in response to 2,3-cGAMP treatment using non-reducing SDS-PAGE. **c,** Cells were pre-treated with NOS inhibitor L-NAME before adding 2,3-cGAMP. STING oligomers were assessed using non-reducing SDS-PAGE. hiMG were serum-starved for 16 h before 2,3-cGAMP treatment for 1 and 6 h. **d,** Ratio of STING oligomers/monomers (70 kDa/35 kDa) after treatment with 2,3-cGAMP normalized to no treatment condition. *n* = 8, Student’s *t* test.

**Extended Data Fig. 2.**
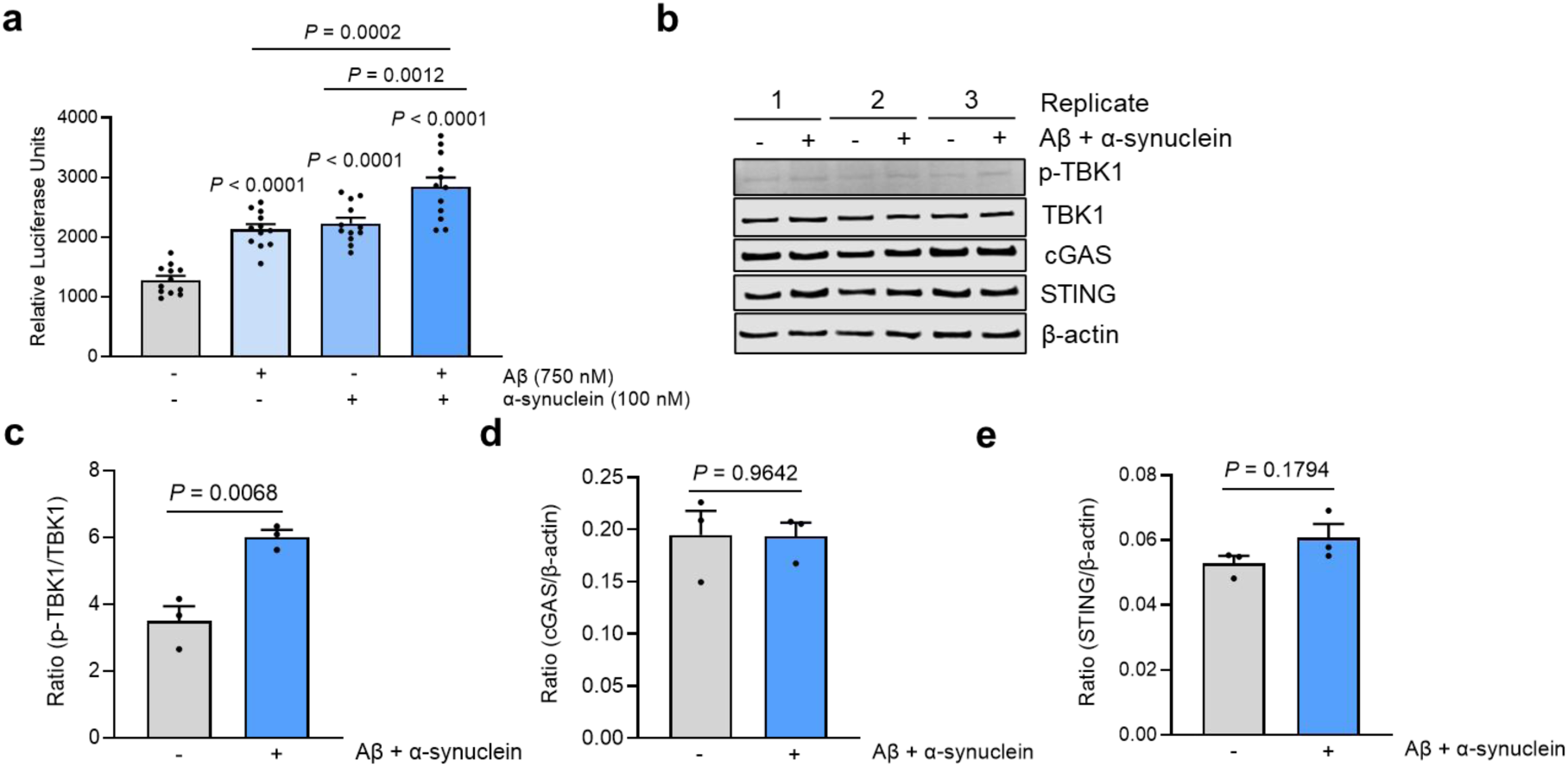
α-Synuclein and Aβ oligomers induce cGAS-STING signaling in hiMG and THP-1 Dual hM. **a,** THP1-Dual WT hM (500,000 cells/well in 1 mL serum-free media in a 12-well plate) were incubated in αSyn oligomers (100 nM), Aβ oligomers (750 nM), or both. Luminescence was monitored at 72 h using freshly prepared QUANTI-Luc reagent (Invivogen). Values indicate mean + s.e.m. (*n* = 12), ANOVA with Tukey’s multiple comparison’s test. **b,** hiMG were incubated with αSyn(100 nM)/Aβ oligomers (750 nM) for 6 h. Cells were lysed, and immunoblotting was performed using p-TBK1, TBK1, cGAS, STING, and β-actin antibodies. **c-e,** Quantification of the ratio of p-TBK1/TBK1, cGAS/β-actin, and STING/ β-actin. Values are mean + s.e.m. (*n* = 3) with Student’s *t* test.

**Extended Data Fig. 3.**
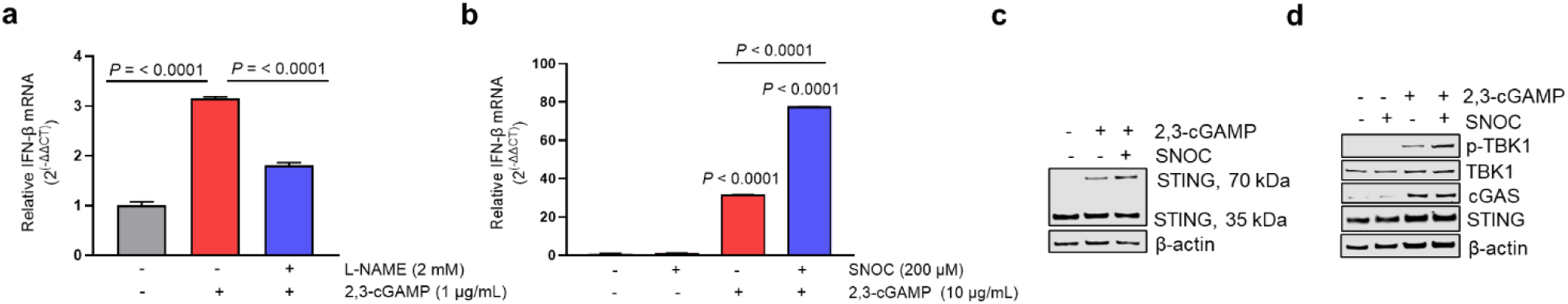
NO-related species boost 2,3-cGAMP-induced cGAS-STING signaling. **a,** Relative human IFN-β mRNA in hM in response to 2,3-cGAMP with and without 3 h pre-treatment with L-NAME. Values are mean + s.e.m., *n* = 3, Student’s *t* test. **b,** Relative IFN-β mRNA expression of mouse BV2 microglia in response to 2,3-cGAMP, SNOC, or both. Cells exposed to SNOC in phosphate buffered saline (PBS, pH 7.4) and 20 min later treated with 2,3-cGAMP in fresh medium for an additional 60 min. Values are mean + s.e.m., *n* = 4, Student’s *t* test. **c,** Non-reducing SDS-PAGE showing STING oligomers and monomers in mouse BV-2 microglia after exposure to SNOC or 2,3-cGAMP. **d,** Reducing SDS-PAGE showing protein levels of p-TBK1, TBK1, cGAS, and STING in mouse BV-2 microglia.

**Extended Data Fig. 4.**
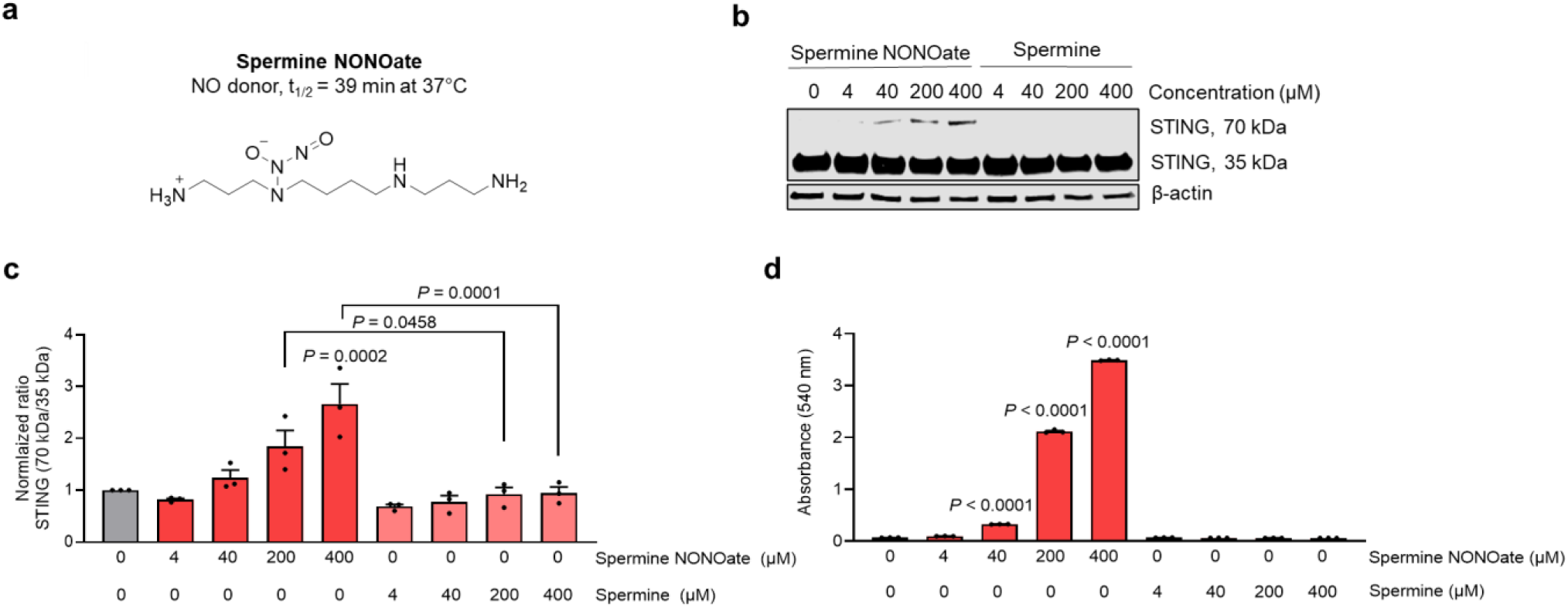
NO donor spermine NONOate induces STING oligomerization. **a,** Chemical structure of spermine NONOate. **b,** Detection of STING monomers and oligomers. hM (2 million cells) were exposed to the indicated concentration of spermine or spermine NONOate for 3 h in phenol red-free medium. Cells were lysed, and non-reducing SDS-PAGE performed, followed by immunoblotting with anti-STING antibody. **c,** Quantification of ratio of STING oligomers/monomers. Values normalized to no treatment and represent the mean + s.e.m. (*n* =3). **d,** Quantification of the stable NO metabolite, nitrite, in media following exposure to spermine NONOate or spermine by Griess assay (Cayman Chemicals). Absorbance at 540 nm reflects the relative nitrite level. Values are mean + s.e.m. (n = 3). Statistics by ANOVA and Tukey’s multiple comparison’s test.

**Extended Data Fig. 5.**
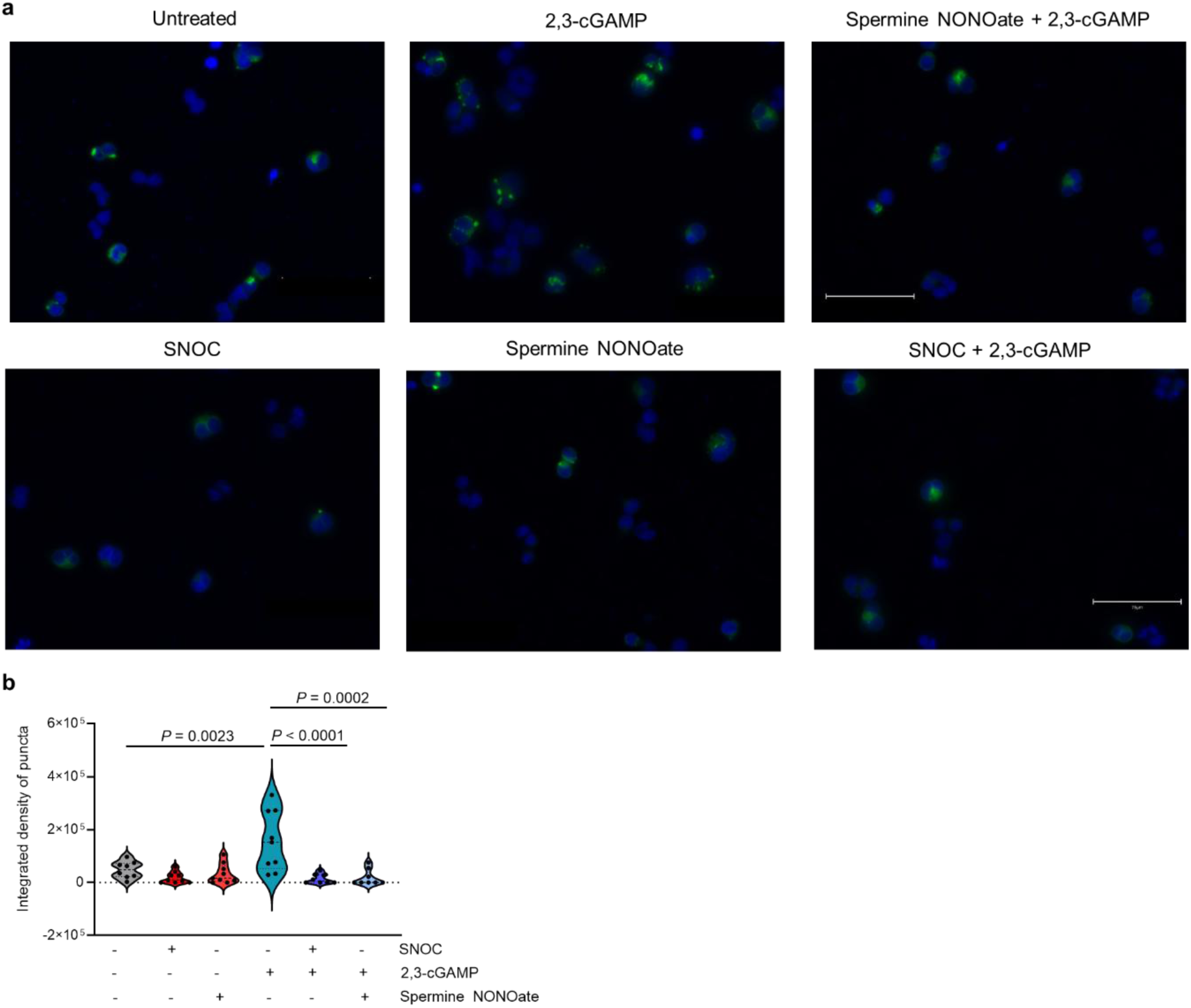
NO Decreases the formation of STING biocondensates. **a,** HEK-293T cells were transfected with STING-EGFP for 18 h, exposed to SNOC (200 μM) or spermine NONOate (400 μM), and analyzed for condensates 1 h later. Scale bars, 75 μm. **b,** Violin plot showing ratio of integrated density of STING-EGFP puncta, representing biocondensates. *n* = 8– 9 for each group with ANOVA and Tukey’s multiple comparison’s test.

**Supplementary Fig. 1.**
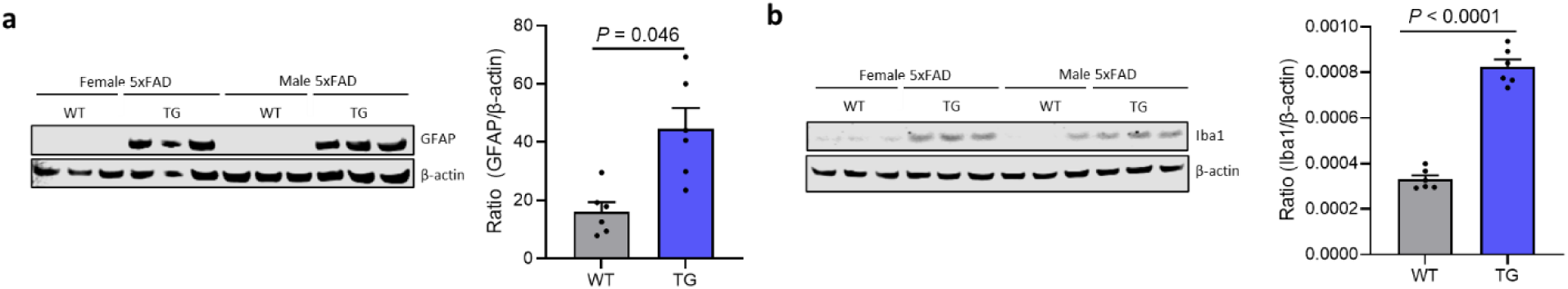
Inflammatory response at 6-months of age in 5xFAD mouse brain. **a, b,** Detection by immunoblot and quantification of GFAP and Iba in 6-month-old 5xFAD TG (*n* = 6) vs, WT (*n* = 6) mice. Values are mean + s.e.m. Statistical significance by unpaired two-tailed Student’s *t* test.

**Extended Data Table 1.**
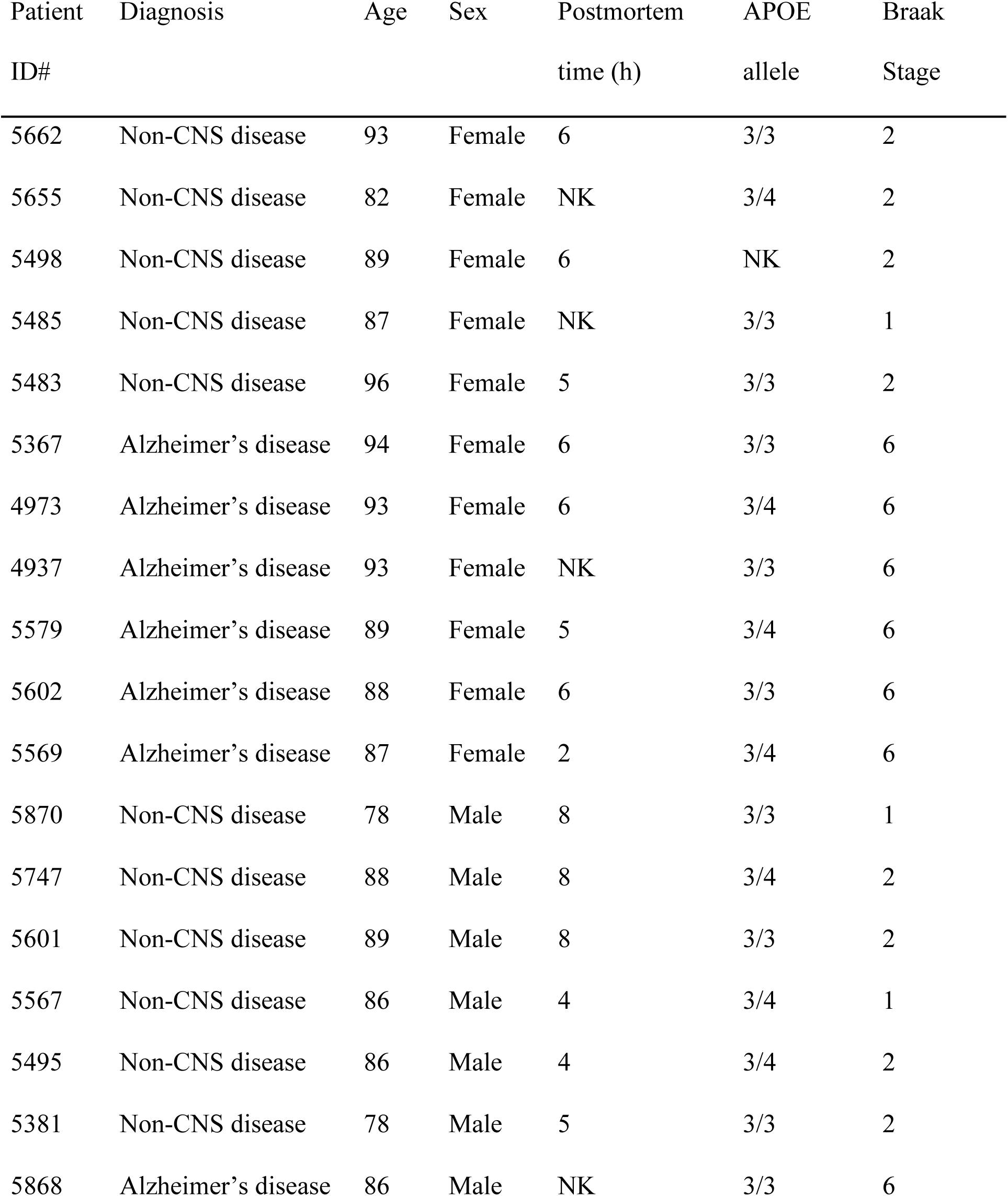
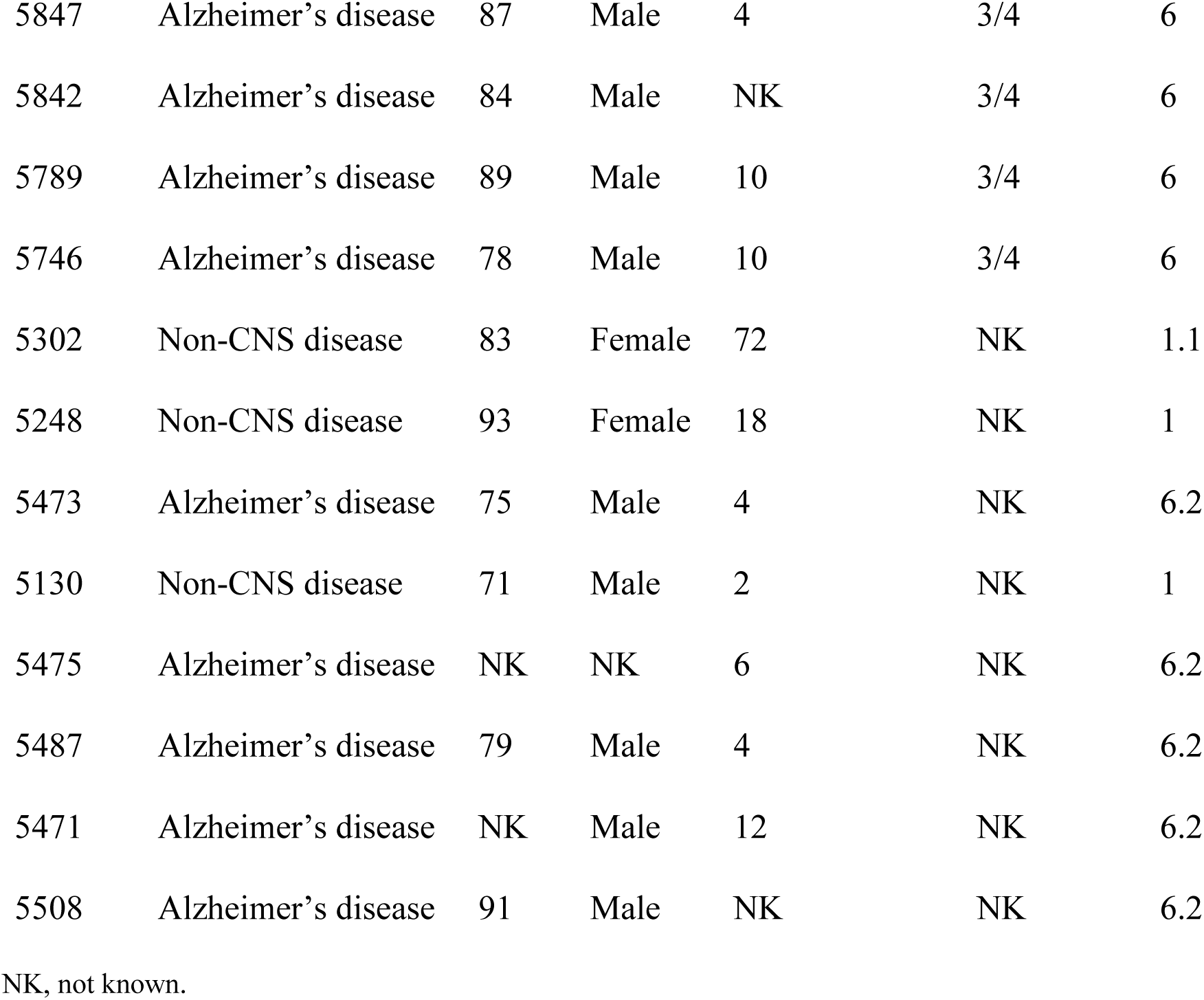
List of human postmortem brains (cerebral cortex) and diagnoses used in this study.

| Patient ID# | Diagnosis | Age | Sex | Postmortem time (h) | APOE allele | Braak Stage |
| --- | --- | --- | --- | --- | --- | --- |
| 5662 | Non-CNS disease | 93 | Female | 6 | 3/3 | 2 |
| 5655 | Non-CNS disease | 82 | Female | NK | 3/4 | 2 |
| 5498 | Non-CNS disease | 89 | Female | 6 | NK | 2 |
| 5485 | Non-CNS disease | 87 | Female | NK | 3/3 | 1 |
| 5483 | Non-CNS disease | 96 | Female | 5 | 3/3 | 2 |
| 5367 | Alzheimer's disease | 94 | Female | 6 | 3/3 | 6 |
| 4973 | Alzheimer's disease | 93 | Female | 6 | 3/4 | 6 |
| 4937 | Alzheimer's disease | 93 | Female | NK | 3/3 | 6 |
| 5579 | Alzheimer's disease | 89 | Female | 5 | 3/4 | 6 |
| 5602 | Alzheimer's disease | 88 | Female | 6 | 3/3 | 6 |
| 5569 | Alzheimer's disease | 87 | Female | 2 | 3/4 | 6 |
| 5870 | Non-CNS disease | 78 | Male | 8 | 3/3 | 1 |
| 5747 | Non-CNS disease | 88 | Male | 8 | 3/4 | 2 |
| 5601 | Non-CNS disease | 89 | Male | 8 | 3/3 | 2 |
| 5567 | Non-CNS disease | 86 | Male | 4 | 3/4 | 1 |
| 5495 | Non-CNS disease | 86 | Male | 4 | 3/4 | 2 |
| 5381 | Non-CNS disease | 78 | Male | 5 | 3/3 | 2 |
| 5868 | Alzheimer's disease | 86 | Male | NK | 3/3 | 6 |
| 5847 | Alzheimer's disease | 87 | Male | 4 | 3/4 | 6 |
| 5842 | Alzheimer's disease | 84 | Male | NK | 3/4 | 6 |
| 5789 | Alzheimer's disease | 89 | Male | 10 | 3/4 | 6 |
| 5746 | Alzheimer's disease | 78 | Male | 10 | 3/4 | 6 |
| 5302 | Non-CNS disease | 83 | Female | 72 | NK | 1.1 |
| 5248 | Non-CNS disease | 93 | Female | 18 | NK | 1 |
| 5473 | Alzheimer's disease | 75 | Male | 4 | NK | 6.2 |
| 5130 | Non-CNS disease | 71 | Male | 2 | NK | 1 |
| 5475 | Alzheimer's disease | NK | NK | 6 | NK | 6.2 |
| 5487 | Alzheimer's disease | 79 | Male | 4 | NK | 6.2 |
| 5471 | Alzheimer's disease | NK | Male | 12 | NK | 6.2 |
| 5508 | Alzheimer's disease | 91 | Male | NK | NK | 6.2 |

NK, not known.

## Online content

Any methods, additional references, Nature Portfolio reporting summaries, source data, extended data, supplementary information, acknowledgements, peer review information; details of author contributions and competing interests; and statements of data are available.

## Methods

### Animal use

All animal studies performed in this work adhered to the NIH Guide for the Care and Use of Laboratory Animals. Animal care was in accordance with institutional guidelines. All human postmortem brains were de-identified for this study and approved for use by the Institutional Review Board. Statistical analyses are included within each figure, when appropriate. Animal studies using immunohistochemistry were blinded during staining, image preparation, imaging, and analysis. Blinding was removed following the final statistical analysis.

### Reagents

Biotin-HPDP as purchased from Dojindo Laboratories (#SB17-10). L-NAME (#N5751 and #80210), ethylenediaminetetraacetic acid (ETDA, #EDS-100G), N-ethylmaleimide (NEM, E3876-5G), L-arginine (#A8094), S-methyl methanethiosulfonate (MMTS, #208795), neocuproine (#N1626), and (+)-sodium L-ascorbate (#11140) were purchased from Sigma Aldrich. Pierce High Capacity NeutrAvidin Agarose (#29204), Dynabeads™ beads A (#10002D), sodium hydroxide (#BP-359-500), TWEEN® 20 (#BP-337-500), NuPAGE™ LDS Sample Buffer 4X (#NP0008), and MES SDS Running Buffer (#B000202) were purchased from Thermo Fisher Scientific. Deoxycholic acid sodium salt (#30970) and Nonidet P 40 substitute (#74385) were purchased from Fluka Biochemika and Fluka Analytical, respectively. Sodium chloride (#7581-06) was purchased from Macron. The following primary antibodies were used: anti-STING (Cell Signaling Technology, 50494; 1/1000 dilution), anti-p-TBK1 (Cell Signaling Technology, 5483S; 1/500), anti-TBK1 (Cell Signaling Technology, 51872S; 1/500), anti-p-IRF3 (Cell Signaling Technology, 4947S; 1/250), anti-IRF3 (Abcam, ab68481, 1/250), anti-cGAS (Santa Cruz, sc-515777; 1/75), and anti-β-actin (Cell Signaling Technology, 3700S; 1/1000). Secondary antibodies were used as follows: IRDye® 800CW-conjugated goat anti-rabbit (1/15,000, Li-Cor, 926-32211) and IRDye® 600VW-conjugated goat-anti mouse (1/15,000, Li-Cor, 926-68020). BSA (bovine serum albumin, #99985) was purchased for primary antibody incubation from Cell Signaling Technologies. Mutant STING constructs were generated using the Q5 ® Site-Directed Mutagenesis Kit (New England BioLabs, E0554S) according to the manufacturer’s protocol. Transformations used BP5α™ Super Competent Cells purchased from BioPioneer Inc (GACC-20). Ambion™ Nuclease-Free Water was purchased from Life Technologies Corp (AM9937). Buffer A contains Hepes-NaOH (100 mM, pH 7.2), EDTA (1 mM), neocuproine (0.1 mM), 1% Nonidet P 40 substitute, 0.4% sodium deoxycholate, 0.075% SDS, and 150 mM NaCl. 2,3-cGAMP was purchased from APExBIO (B8362) or InvivoGen (tlrl-nacga23s). Experiments were performed using 2,3-cGAMP from APExBIO unless otherwise specified.

### Cell line cultures

Human THP-1 monocyte cells (hM) were obtained from ATCC (TIB-202). hM were cultured using phenol-red free RMPI (Quality Biological #112-040-101) containing GlutaMax (2 mM), 10% FBS, and a mixture of 100 U/ml penicillin-100 μg/ml streptomycin (Pen-Strep), and experiments were performed without serum. hM Dual cells were obtained from InvivoGen (thpd-nfis). hM Dual cells were cultured in RPMI 1640 containing 25 mM HEPES, 10% heat-inactivated fetal bovine serum,100 μg/ml Normocin^™^, Pen-Strep (100 U/ml-100 μg/ml), Blasticidin (10 mg/ml), and Zeocin^®^ (100 mg/ml), and experiments were performed without serum. BV2 mouse microglia were obtained from Accegen (ABC-TC212S) cultured using DMEM (Gibco # 10-566-016) containing 10% FBS and 1% pen/strep.

### Human induced pluripotent stem cell (hiPSC)–derived microglia

hiPSCs were generated from human fibroblasts (Hs27, ATCC, CRL-1634) and cultured as previously described.^12,29^ hiPSC-derived microglia (hiMG) were differentiated using our previously published protocol.^12,29^ hiMG were placed in phenol red-free media (Iscove’s modified Dulbecco’s medium (IMDM; Thermo Fisher Scientific, 21056023) the day after the cells were plated and 16 h before the start of each experiment.

### Synthesis of *S*-nitrosocysteine (SNOC)

As described.^52^ 100 mM S-nitrosocysteine (SNOC) stock solution was synthesized immediately before use, and all steps were performed in the dark. Briefly, L-cysteine (15 mg, Sigma Aldrich, #30089) was dissolved in 588.8 μL sterile distilled water (Gibco, #15230170), and NaNO_2_ (10 mg, Sigma Aldrich, #237213) was dissolved in a separate tube containing 688.4 μL sterile distilled water. Both solutions were placed on ice, and 475 μL of each solution was mixed into a new tube. This new solution was kept on ice and in the dark for 5 min; 50 μL 10 M HCl was added, and a working SNOC solution was made using PBS pH 7.4 (Gibco, #10010023). SNOC was used immediately after adding HCl and was kept on ice throughout the process. All experimental exposures to SNOC were performed at room temperature, in the dark, and away from direct sunlight and/or UV lamps.

### Detection of *S-*nitrosylated proteins by biotin-switch assay

The biotin-switch assay procedure was based on a previously described protocol.^53^ It is important to note that all steps before and after incubating the biotinylated proteins with agarose beads were performed in the dark and away from direct sunlight. After cells were stimulated, they were harvested and lysed in 400 μL ice-cold Buffer A containing 10 mM S-methyl methanethiosulfonate (MMTS) on ice. Buffer A contained Hepes-NaOH pH 7.2 (100 mM), EDTA (1 mM), neocuproine (0.1 mM), 1% Nonidet P 40 substitute, 0.4% sodium deoxycholate, 0.075% SDS, and 150 mM NaCl. Lysed cells were immediately centrifuged at 17,799 x g at 4 °C, and the supernatant was collected. The supernatant was placed in a new tube containing 32 μL 25% SDS and was then shaken on a heat block at 45 °C for 20 min at 850 rpm using an Eppendorf Thermomixer. Two volumes of 100% ice-cold acetone (stored at −20 °C) were added to each sample to precipitate proteins and remove excess MMTS. The samples were vigorously shaken six times before incubating at −20°C for 20 min. Samples were then centrifuged at 3266 x g for 5 min at 4 °C and the supernatant was removed. An aliquot of 700 μL of ice-cold 70% acetone (30% water and 70% acetone, stored at −20°C) was added to each sample on ice.

Samples were briefly vortexed (3 s) and then centrifuged at 3266 x g for 5 min at 4°C. The supernatant was removed, and samples were centrifuged for an additional 1 min to remove any remaining acetone. The pellet was resuspended on ice in 100 μL Hepes-NaOH pH 7.2 (100 mM) containing EDTA (1 mM), neocuproine (0.1 mM), and 1% SDS. Biotin-HPDP (5 μL) was added to each sample, followed by the addition of 10 μL of a 200 mM (+)-sodium (L)-ascorbate solution. The 200 mM (+)-sodium (L)-ascorbate solution was prepared immediately before use by solubilizing 19.8 mg in 500 μL Hepes-NaOH pH 7.2 (100 mM) containing EDTA (1 mM) and neocuproine (0.1 mM), and stored on ice in the dark. Negative controls did not have (+)-sodium (L)-ascorbate added. Samples were gently mixed by flicking the tube and incubated in the dark at room temperature for 1 h. During this time, Neutravidin agarose beads (30 μL/sample + one extra) were washed twice using 1 mL PBS pH 7.4 by gently spinning down beads and removing the supernatant. After the beads were washed, they were resuspended in PBS 7.4 (100 μL/sample + one extra) and transferred to new tubes on ice. After the 1 h incubation at room temperature, 600 μL 100% ice-cold acetone (stored at −20°C) was added to each sample.

Samples were vigorously shaken six times before incubating at −20 °C for 20 min. Samples were then centrifuged at 3266 x g for 5 min at 4 °C and the supernatant was removed. 600 μL of ice-cold 70% acetone (30% water and 70% acetone, stored at −20°C) was added to each sample on ice. Samples were briefly vortexed (3 s) and then centrifuged at 3266 x g for 5 min at 4 °C. The supernatant was removed, and samples were centrifuged for an additional 1 min to remove any remaining acetone. After acetone removal, the pellet was resuspended on ice in 100 μL Hepes-NaOH pH 7.2 (100 mM) containing EDTA (1 mM), neocuproine (0.1 mM), and 1% SDS. Two volumes of PBS pH 7.4 was added to each sample, and samples were centrifuged at 12,175 x g for 1 min at 4 °C. 15 μL of the supernatant was saved for the “input,” and the remaining volume was added to Neutravidin agarose beads (30 μL beads/sample). Agarose beads were allowed to tumble overnight with biotinylated protein samples at 4°C. LDS buffer (90 μL 4X LDS + 10 μL β-mercaptoethanol, 5 μL/sample) was added to each of the input samples. Reducing input samples were boiled for 3 min and then stored at −80 °C until all samples were ready to be run with SDS-PAGE. The next day, samples were washed twice with 500 μL Hepes-NaOH pH 7.2 (100 mM) containing EDTA (1 mM), neocuproine (0.1 mM), and 1% SDS by centrifuging at 2516 x g for 1 min at room temperature. Samples were washed with 150 μL PBS pH 7.4 and centrifuged at 2516 x g for 1 min at room temperature. LDS buffer (90 μL 4X LDS + 10 μL β-mercaptoethanol, 7.5 μL/sample) was added to each sample, and samples were boiled for 5 min. SDS-PAGE, followed by immunoblotting, was performed to visualize *S*-nitrosylated proteins and input protein levels. Immunoblotting was performed using Bolt™ 4-12%, Bis-Tris Plus WedgeWell™ gels, MES SDS running buffer, and Bolt™ transfer buffer. PVDF membranes were incubated with 10 mL Intercept (TBS) blocking buffer (LiCOR) at room temperature for 1 h. Primary antibodies were incubated in TBST (1X) containing 0.05% BSA overnight at 4 °C. PVDF membranes were briefly washed 3 times with TBST (1X). PVDF membranes were then incubated in 5 mL TBST (1X) and 5 mL Intercept (TBS) blocking buffer (LiCOR) (1:1 ratio) containing 0.1% SDS, 0.02% Tween-20, and secondary antibodies (LiCOR). The signal was detected using an Odyssey CLx Li-COR instrument. If the negative control for the biotin-switch assay failed in a sample preparation, then that sample was omitted from the analysis.

### Detection of SNO-STING in hM

One million hM/mL were plated (2 mL) in a 6-well plate. After 24 h, cells were centrifuged to obtain a cell pellet. The cell pellet was either exposed to PBS pH 7.4 alone, PBS pH 7.4 containing 200 μM SNOC, or old SNOC. After 20 min, the cells were centrifuged, and PBS was removed. Cells were resuspended in Buffer A (Hepes-NaOH pH 7.2 (100 mM), EDTA (1 mM), neocuproine (0.1 mM), 1% Nonidet P 40 substitute, 0.4% sodium deoxycholate, 0.075% SDS, and 150 mM NaCl) containing 10 mM MMTS. The biotin-switch assay was performed on the lysate as previously described.^53^ Ascorbic acid was not added to one of the SNOC-treated samples as a negative control. *S*-Nitrosylated STING was detected following SDS-PAGE and western blotting using a primary STING antibody (1:1000, Cell Signaling #50494).

### Detection of SNO-STING in hiMG

Eight million hiMG were used for each treatment condition. Culture medium was removed, and cells were either exposed to PBS pH 7.4 alone, PBS pH 7.4 containing 200 μM SNOC, or old SNOC. After 20 min, the cells were centrifuged, PBS was removed, and a cell lysate was prepared and analyzed by biotin-switch assay as above.

### Detection of 2,3-cGAMP-induced SNO-STING in hM

One million hM/mL were plated (2 mL) in a 6-well plate. Cells were serum-starved 24 h before the experiment. After 24 h, cells were pre-treated with L-NAME (2 mM) for 3 h and then treated with 2,3-cGAMP (10 μg/mL). Experiments were performed using 2,3-cGAMP from APExBIO unless specified. After 24 h, cells were centrifuged to obtain a cell pellet. The cell pellet was resuspended to produce a cell lysate and analyzed by biotin-switch assay as above.

### Detection of Aβ/αSyn oligomer-induced SNO-STING in hiMG

Eight million hiMG were used for each condition. Cells were serum-starved for 16 h before adding L-NAME (2 mM). L-Arginine levels were brought to 1 mM in all samples. Oligomers (Aβ, 750 nM; αSyn, 150 nM) were added 3 h after a 24-treatment with L-NAME (2 mM) and were prepared according to the previous protocol.^29^ Medium was removed, and cells were resuspended to produce a cell lysate and analyzed by biotin-switch assay as above.

### Detection of SNO-STING in human postmortem brain tissue

Postmortem cerebrocortical tissue from AD patients and patients dying of non-CNS causes was lysed in Buffer A containing protease/phosphatase inhibitor (Cell Signaling, 5872S) using a Dounce homogenizer (1 mL). The proteome was normalized to 1.5 mg, and the total volume (200 μL) was adjusted using Hepes-NaOH pH 7.2 (100 mM) containing EDTA (1 mM) and neocuproine (0.1 mM). Biotin-switch assays were performed as above.

### Generation of non-nitrosylatable mutant STING constructs

Mutant non-nitrosylatable STING constructs were generated using the Q5^®^ Site-Directed Mutagenesis Kit (New England BioLabs, E0554S) according to the manufacturer’s protocol. Transformations used BP5α™ Super Competent Cells purchased from BioPioneer Inc (GACC-20).

### Identification of Critical SNO-Cysteines using Mass Spectrometry

Full-length human recombinant STING protein (10 µg, Origene, TP308418, lot: RK035D61) was added to PBS pH 7.4 on ice in the dark. Samples were incubated with or without 10 μM or 25 μM freshly prepared SNOC at room temperature in the dark. After 20 min, ice-cold buffer A containing MMTS was added to each sample to a final concentration of 10 mM MMTS. SDS (2%) was added to each sample. Samples were then shaken at 850 rpm in an Eppendorf Thermomixer for 20 min at 45 °C. Next, 2 × volumes of ice-cold 100% acetone was added to each sample and vigorously shaken six times before incubating at −20°C for 20 min. Samples were then centrifuged at 3266 x g for 5 min at 4 °C and the supernatant was removed. The pellet was resuspended in 180 μL Hepes-NaOH pH 7.2 (100 mM) containing EDTA (1 mM), neocuproine (0.1 mM), and 1% SDS.

Next, for analysis by mass spectrometry, biotin-switch chemistry using NEM instead of biotin was performed in order to label free cysteine residues that had been *S*-nitrosylated. Accordingly, samples were incubated with or without 20 μL 200 mM sodium-L-ascorbate and 2 μL of a 100 mM NEM stock for 1 h at room temperature in the dark. At the final resuspension step, 100 μL of PBS pH 7.4 was added, and samples were frozen at −80 °C. Samples were subsequently thawed and precipitated by methanol/chloroform, and then redissolved in 8 M urea/100 mM TEAB, pH 8.5. Proteins were reduced with 5 mM tris(2-carboxyethyl)phosphine hydrochloride (TCEP, Sigma-Aldrich) and alkylated with 10 mM chloroacetamide (Sigma-Aldrich). Proteins were digested overnight at 37 °C in 2 M urea/100 mM TEAB, pH 8.5, with trypsin (Promega). Digestion was quenched with formic acid, 5% final concentration. The digested samples were analyzed on an Orbitrap Eclipse tribrid mass spectrometer (Thermo Scientific). The digest was injected directly onto a 25 cm, 100 um ID column packed with BEH 1.7um C18 resin (Waters). Samples were separated at a flow rate of 300 nL/min on a nLC 1200 (Thermo). Buffer A and B were 0.1% formic acid in 5% and 90% acetonitrile, respectively. A gradient of 0-25% B over 100 min, an increase to 40% B over 20 min, an increase to 90% B over 10 min and held at 90%B for a final 10 min was used for 140 min total run time. Column was re-equilibrated with 15 µL of buffer A prior to the injection of sample. Peptides were eluted directly from the tip of the column and nanosprayed directly into the mass spectrometer by application of 2.5 kV voltage at the back of the column. The Eclipse was operated in data-dependent mode. Full MS scans were collected in the Orbitrap at 120k resolution with a mass range of 375 to 1500 m/z and an AGC target of 4e^5^. The cycle time was set to 3 s, and within this 3 s the most abundant ions per scan were selected for CID MS/MS in the ion trap. Quadrupole isolation at 1.6 m/z was used, monoisotopic precursor selection was enabled, and dynamic exclusion was used with exclusion duration of 10 s.

Protein and peptide identification were performed with Integrated Proteomics Pipeline – IP2 (Integrated Proteomics Applications). Tandem mass spectra were extracted from raw files using RawConverter^54^ and searched with ProLuCID^55^ against the Uniprot human database. The search space included all fully-tryptic and half-tryptic peptide candidates. Carbamidomethylation and NEM on cysteine were considered as a differential modifications. Data were searched with 50 ppm precursor ion tolerance and 600 ppm fragment ion tolerance. Identified proteins were filtered to using DTASelect^56^ and utilizing a target-decoy database search strategy to control the false discovery rate to 1% at the protein level.^57^

### Detection of cGAS-STING protein markers in hiMG

hiMG were lysed in M-PER™ (Thermo Scientific, 78501) containing protease/phosphatase inhibitor (Cell Signaling, 5872S). Lysates were shaken at 850 rpm in an Eppendorf Thermomixer at 4 °C for 20 min, followed by centrifugation at 12,298 x g for 5 min at 4°C. The proteome was normalized in all samples, and LDS buffer samples were made. For chemical reduction, LDS samples contained 6% β-mercaptoethanol and were boiled for 5 min before performing SDS-PAGE and immunoblotting. Non-reducing samples did not contain β-mercaptoethanol and were not boiled.

### Detection of cGAS-STING pathway proteins in human postmortem brain tissue

Cerebrocortical tissue was lysed in Buffer A containing protease/phosphatase inhibitor (Cell Signaling, 5872S) using a Dounce homogenizer (1 mL). Lysates were shaken at 850 rpm using an Eppendorf Thermomixer at 4 °C for 20 min, followed by centrifugation at 12,298 x g for 5 min at 4 °C. The proteome was normalized in all samples, and LDS buffer samples were made. Samples contained 6% β-mercaptoethanol and were boiled for 5 min before performing SDS-PAGE and immunoblotting.

### STING oligomerization time course experiment

hM cells were plated in a 6-well plate containing 2 mL serum-free DMEM medium containing 1% pen/strep. After 3 h, cells were treated with 10 μg/mL 2,3-cGAMP. The cell pellet was obtained at each time point and resuspended in Buffer A containing protease/phosphatase inhibitors (Cell Signaling, 5872S). Samples were shaken at 850 rpm in an Eppendorf Thermomixer at 4 °C for 20 min and then centrifuged at 12,298 x g for 5 min at 4 °C. The proteome was quantified, and a non-reducing SDS-PAGE gel was run, followed by western blotting. STING monomers and oligomers were visualized with a primary STING antibody (1:1000, Cell Signaling #50494).

### STING oligomerization using human full-length recombinant STING

Recombinant STING protein (0.5 µg, Origene, TP308418, lot: RK029BS) was added to an Eppendorf tube containing PBS pH 7.4. Then, 0, 25, 50, 100, or 200 μM SNOC was added to each tube. After 20 min, an equal volume 2x LDS buffer without reducing agent was added immediately to each sample. Non-reducing SDS-PAGE was performed, followed by western blotting. STING monomers and oligomers were visualized as above.

### Concentration-dependent SNO-STING formation

hM (500,000/mL) were plated in 2 mL RPMI containing 10% FBS and 1% Pen-strep. After 16 h, cells were centrifuged, and medium was removed. Cells were treated with either PBS pH 7.4 (alone) or PBS pH 7.4 containing SNOC at 25, 50, 100, or 200 μM at room temperature. After 20 min, cells were centrifuged, the supernatant removed, and resuspended in Buffer A. Biotin-switch assays were then performed. Non-reducing and reducing SDS-PAGE were also performed, followed by western blotting. STING monomers, oligomers, and SNO-STING were visualized as above.

### Formation of STING oligomers after spermine NONOate exposure

hM (1 million/mL) were plated in 2 mL RPMI containing 10% FBS and 1% Pen-strep. Cells were serum-starved 24 h before compound treatment. Spermine NONOate (or spermine control) was solubilized in 0.01 M NaOH, kept on ice, protected from light, and used immediately after solubilization. Cells were treated with equal concentration and volume of either spermine or spermine NONOate for 3 h at 37 °C. Cells were then centrifuged and lysed on ice using Buffer A containing 100x protease/phosphatase inhibitors. Lysates were shaken at 850 rpm in an Eppendorf Thermomixer for 20 min at 4 °C. Lysates were then centrifuged at 12,298 x g for 5 min at 4 °C. The proteome was normalized, and a non-reducing SDS-PAGE was performed. Immunoblotting was performed, and STING antibody was used to detect oligomers and monomers.

### STING condensate formation

HEK 293T cells (20,000/mL) were plated in serum-free DMEM media with 1% Pen/Strep on a 24-well plate. Fugene-HD was used to transfect human STING-EGFP(C-terminal tag). Briefly, 2 μg plasmid DNA was added to 100 μL OptiMEM media with 3 μL Fugene-HD. This solution was incubated at room temperature for 20 min. HEK 293T cells were plated during this incubation. The DNA-Fugene mixture (25 μL)was added to each well after the cells were plated. Cells were treated with 10 μg/mL 2,3-cGAMP and/or freshly prepared SNOC (200 μM) or spermine NONOate (400 μM) at 18 h-post transfection. After 1 hr, cells were imaged using an EVOS microscope at 40X. Hoechst was added to cells at a final concentration of 1 μg/mL after the treatment was completed. Hoechst was incubated with the cells for 10 minutes before imaging at 37 °C. Scale bars, 75 μm. Puncta were quantified using ImageJ according to the previous publication with the following modifications. The rectangle tool was used instead of the polygon tool; 49 and 255 were used for the lower and higher limit for the threshold, and 1-200 was used as the particle size.

### THP-1 Dual cell IRF luciferase signaling assay

Aliquots of 100,000 THP-1 Dual cells in 100 μL serum-free RPMI Dual medium containing a stable interferon (IRF) luciferase reporter were treated with various stimuli. After incubation for the indicated time point, a 20 μL aliquot of cell medium from each experimental sample was placed in a white bottom 96-well plate. Then, 50 μL of freshly prepared Quanti-LUC (Invivogen, #rep-qlc4lg1) was added to each well immediately before quantifying the luciferase signal using a plate reader (Molecular Devices, SpectaMax M3). An integration time of 1000 ms was used.

### Griess assay

A Griess assay was performed to measure nitrite levels according to the manufacturer’s instructions (Cayman Chemical, #780001). Assays were performed on 100 μL aliquots of medium.

### Immunohistochemistry

Mice were anesthetized with isoflurane (1% - 2%) and the withdrawal reflex was checked to assure adequate depth of anesthesia. An incision was made to reveal the thoracic cavity, and the heart was exposed. A 23-gauge sterile butterfly catheter was inserted through the left ventricle into the ascending aorta, and the right atrium was sniped. After sacrifice, all animals were perfused transcardially with ice-cold PBS followed by 4% paraformaldehyde (PFA) for fixation. After perfusion was complete, the animals were decapitated and the brains were isolated quickly using a caudal approach. The isolated brains were incubated in 4% PFA overnight at 4°C for post-fixation. The following day, brains were rinsed with PBS and incubated in 30% sucrose at 4 °C until sectioning. All brain identities were blinded until the completion of all the analyses. Approximately 40 μm-thick sagittal sections were made with a vibratome (Leica VT1200S) and incubated in blocking buffer (5% normal goat serum, 0.5% Triton X-100 in PBS) at room temperature for one hour to prevent non-specific binding. Following the blocking step, all floating sections were incubated overnight at 4 °C on a shaker in primary antibody solutions containing one of the following: Ionized calcium binding adaptor molecule 1 (Iba1) (EPR16589, 1:1000, #ab178847, Abcam), Synapsin I (D12G5, 1:500, rabbit mAb #5297, Cell Signaling Technology), anti-RFP (1:500, #600-401-379, Rockland). The following day, sections were rinsed with PBS and transferred to their respective secondary antibody solutions for 2 hours at room temperature on a shaker: Alexa 488 (1:500, goat anti-Mouse, A11001, Life Technologies), Alexa 488 (1:500, goat anti-rabbit, A11008, Life Technologies), Alexa 568 (Goat anti-rabbit, A11011, Life Technologies). The sections were rinsed again with PBS, mounted on Superfrost plus slides (12-550-15, Fisher scientific) and dried at room temperature. The slides were coated with Vectashield plus antifade mounting medium with DAPI (H-2000, Vector Laboratories), and coverslip (12541057, Fisher Scientific) was applied. Images were acquired with a 20x magnification lens at Zoom 2 using a confocal microscope (Nikon Eclipse Ti2). Sections that showed weak to no tdTomato expression, or those that were over-saturated were excluded from the analysis as part of our quality control procedures. Post-image processing and quantifications were performed using Fiji ImageJ (1.54f).^58^ For Iba1 quantification, uniform sized ROIs were manually placed over areas containing Iba1 (green) and tdTomato (red), and Iba1 integrated density was calculated and compared. For Synapsin I quantification, ROIs were manually placed around the tdTomato-positive region (red), and Synapsin I integrated density was calculated. The integrated density was normalized to the area of the ROI for final quantification and then compared. As part of our data quality control, an outlier test (ROUT method) was performed, and a few rare outliers were excluded from the analysis.

### Quantitative polymerase chain reaction (qPCR)

hM and BV-2 cells were treated with 2,3-cGAMP for 24 h prior to RNA isolation. RNA was isolated using the Quick RNA Miniprep kit (Zymo Research Corporation, 50-444-597) and reverse transcription was performed according to the QuantiTect Reverse Transcription βkit (Qiagen, 205311) instructions. qPCR was then performed using SYBR reagents (Quantabio, 101414-270) to quantify the relative IFN-β mRNA levels. The cycle threshold (Ct) values for IFN-β were normalized to the 18s housekeeping gene control, and relative gene expression was calculated using the relative quantification method (2^−ΔΔCt^). Ambion™ Nuclease-Free Water was purchased from Life Technologies Corp (AM9937). The 18s primers used were 5’ GTCGTAGTTCCGACCATA 3’ (Forward, human), 5’ GCCCTTCCGTCAATTCCTTT 3’ (Reverse, human), 5’ GGACCAGAGCGAAAGCATTTGCC 3’ (Forward, mouse), and 5’ TCAATCTCGGGTGGCTGAACGC 3’. The IFN-β primers used were 5’ GCTTGGATTCCTACAAAGAAGCA 3’ (Forward, human), 5’ ATAGATGGTCAATGCGGCGTC 3’ (Reverse, human), 5’ TCCGAGCAGAGATCTTCAGGA 3’ (Forward, mouse), and 5’ TGCAACCACCACTCATTCTGAG 3’ (Reverse, mouse).

## Acknowledgements

This work was supported in part by NIH grants R35 AG071734, U01 AG088679, RF1 AG057409, R01 AG078756, R01 AG056259, R01 DA048882 and DP1 DA041722 (to S.A.L.), and R01 AG077046 (to J.R.Y. III). Additional support was from DoD/Department of the Army Grant AR230101 (to S.A.L.), and NIA/NIH HOPE training grant T32 AI007384 and Multidisciplinary Training in Basic and Translational Alzheimer’s Disease Research training grant T32 AG066596-05 fellowship awards (to L.N.C.).

## Author contributions

S.A.L. conceived of the study. L.N.C. designed the study, performed biochemical, cell- and animal-based experiments, performed experiments with human postmortem patient samples, performed imaging and immunohistochemistry experiments, and analyzed results. P.B. performed immunohistochemistry and the corresponding analysis. L.N.C., X.Z., T.N., and S.A.L coordinated the research to obtain brain tissues from mice and the UCSD Brain Bank. A.R. performed lentiviral injections related to immunohistochemistry. J.N. and E.S. prepared hiPSC-derived microglia (hiMG), and E.S. performed imaging of the hiMG. J.D. performed the mass spectrometry experiment and interpreted results with J.R.Y. III overseeing this work. X.Z. performed preliminary experiments showing protein *S*-nitrosylation of STING. C.R., P.P., H.S., and N.L. isolated brain samples from mice. L.N.C. and S.A.L wrote the manuscript.

## Competing interests

S.A.L. discloses that he is an inventor on worldwide patents for the use of memantine and NitroSynapsin for neurodegenerative and neurodevelopmental disorders. Per Harvard University guidelines, he participates in a royalty-sharing agreement with his former institution, Boston Children’s Hospital/Harvard Medical School, which licensed the FDA-approved drug memantine (Namenda®) to Forest Laboratories, Inc./Actavis/Allergan/AbbVie. S.A.L. is also a scientific founder of Adamas Pharmaceuticals, Inc. (now owned by Supernus Pharmaceuticals, Inc.), which developed or comarkets FDA-approved forms of memantine- or amantadine-containing drugs (NamendaXR^®^, Namzaric^®^, and GoCovri^®^), and of EuMentis Therapeutics, Inc., which has licensed NitroSynapsin and related aminoadamantane nitrates from S.A.L. He also serves on the Scientific Advisory Board of Point6 Bio. T.N. serves as a consultant to Dojindo Molecular Technologies. The other authors declare no competing interests.

## Notes

### Competing Interest Statement

SAL discloses that he is an inventor on worldwide patents for the use of memantine and NitroSynapsin for neurodegenerative and neurodevelopmental disorders. Per Harvard University guidelines he participates in a royalty-sharing agreement with his former institution Boston Childrens Hospital/Harvard Medical School which licensed the FDA-approved drug memantine (Namenda) to Forest Laboratories, Inc./Actavis/Allergan/AbbVie. S.A.L. is also a scientific founder of Adamas Pharmaceuticals Inc. (now owned by Supernus Pharmaceuticals Inc.) which developed or comarkets FDA-approved forms of memantine- or amantadine-containing drugs (NamendaXR, Namzaric, and GoCovri) and of EuMentis Therapeutics, Inc. which has licensed NitroSynapsin and related aminoadamantane nitrates from S.A.L. He also serves on the Scientific Advisory Board of Point6 Bio. T.N. serves as a consultant to Dojindo Molecular Technologies. The other authors declare no competing interests.

